# Friendly regulates membrane depolarization induced mitophagy in Arabidopsis

**DOI:** 10.1101/2020.07.12.198424

**Authors:** Juncai Ma, Zizhen Liang, Jierui Zhao, Pengfei Wang, Wenlong Ma, Juan A. Fernandez Andrade, Yonglun Zeng, Nenad Grujic, Liwen Jiang, Yasin Dagdas, Byung-Ho Kang

## Abstract

The oxidative environment within the mitochondria makes them particularly vulnerable to proteotoxic stress. To maintain a healthy mitochondrial network, eukaryotes have evolved multi-tiered quality control pathways. If the stress cannot be alleviated, defective mitochondria are selectively removed by autophagy via a process termed mitophagy. Despite significant advances in metazoans and yeast, in plants, the molecular underpinnings of mitophagy are largely unknown. Here, using time-lapse imaging, electron tomography and biochemical assays, we show that uncoupler treatments cause loss of mitochondrial membrane potential and induce autophagy in Arabidopsis. The damaged mitochondria are selectively engulfed by autophagosomes that are ATG5 dependent and labelled by ATG8 proteins. Friendly, a member of the Clustered Mitochondria protein family, is recruited to the damaged mitochondria to mediate mitophagy. In addition to stress, mitophagy is also induced during de-etiolation, a major cellular transformation during photomorphogenesis that involves chloroplast biogenesis. De-etiolation triggered mitophagy regulates cotyledon greening, pointing towards an inter-organellar cross-talk mechanism. Altogether our results demonstrate how plants employ mitophagy to recycle damaged mitochondria during stress and development.

## Introduction

Mitochondria are highly dynamic double-membraned organelles that function as cellular powerhouses. They generate energy via oxidative phosphorylation (OXPHOS) and mediate the synthesis of essential macromolecules such as iron-sulfur clusters^1,2^. One of the by-products of the oxidative environment in mitochondria is generation of toxic reactive oxygen species that damage mitochondrial DNA, lipids and proteins. In addition, although most of the mitochondrial proteins are encoded by nuclear genes, 13 subunits of the oxidative phosphorylation complexes are still encoded by the mitochondrial genome. As the inter-genome coordination could be disrupted, and one cell could have thousands of times more copies of mitochondrial genome than the nuclear genome; imbalances in stoichiometries of these multi-subunit OXPHOS complexes trigger proteotoxic stress^3,4^. To overcome these challenges and maintain a healthy mitochondrial network, eukaryotes have evolved multi-tiered and interconnected mitochondrial quality control pathways^3^.

One of the major mitochondrial quality control pathways is mitophagy, the selective removal of damaged or superfluous mitochondria via autophagy. As many players involved in mitophagy have been associated with disease, and mitophagy allows us to visualize selective engulfment of an organelle into an autophagosome, mitophagy is one of the best studied signalling mechanisms in metazoans^5–8^. One of the hallmarks of damaged mitochondria is loss of membrane potential^3^. Various chemical protonophores such as carbonyl cyanide p-trifluoro-methoxyphenyl hydrazone (FCCP) or 2,4-dinitrophenol (DNP) have been used to induce mitochondrial membrane depolarization and mitophagy^9^. Loss of mitochondrial membrane potential leads to the stabilization of PINK1 on mitochondrial outer membrane. PINK1 phosphorylates ubiquitin and activates Parkin on mitochondrial membrane for polyubiquitination of various outer membrane proteins. This creates a positive feedback loop that results in recruitment of various selective autophagy receptors such as p62, Optineurin or NDP52 to recruit the damaged mitochondria into autophagosomes for their subsequent degradation^7,10,11^. Although much has been learnt about mitophagy in metazoans, molecular players that mediate mitophagy in plants is currently unknown^2,12^.

Mounting evidence suggests plant mitochondria are also recycled by selective autophagy^2,13^. However, likely influenced by harbouring another endosymbiotic organelle, plants lack homologs of known mitophagy receptors and regulators^12^. Also, so far, most studies used genetic and biochemical assays to analyse mitochondrial turnover in plants. Cell biological tools that would allow us to visualize different stages of mitophagy have not been established. Here, we studied uncoupler induced mitophagy in the model plant *Arabidopsis thaliana*. We used live cell imaging and electron tomography to visualize the engulfment of the damaged mitochondria by mitophagosomes. We supported our cell biological findings with autophagic flux assays to show autophagy regulates recycling of damaged mitochondria. We also showed that Friendly (FMT) protein that has been linked to the regulation of mitochondria dynamics is recruited to mitochondria upon damage. Consistently, *fmt* mutants have defects in formation of mitophagosomes and mitochondrial turnover. Finally, we demonstrate that de-etiolation also leads to accumulation of compromised mitochondria and induces mitophagy. Altogether, our findings establish a cell biological and biochemical platform to further dissect mitophagy and reveal a molecular player that is essential for mitophagy in plants.

## Results

### Uncoupler treatments induce accumulation of depolarized mitochondria in Arabidopsis root cells

Uncouplers such as DNP and FCCP perturb the electrochemical potential of inner mitochondrial membrane, triggering mitochondrial recycling in mammalian cells^14^. To test whether these compounds also affect mitochondria in plant root tip cells, we incubated *Arabidopsis* seedlings expressing mitochondrion-targeted GFP (Mito-GFP) in liquid MS medium containing DNP or FCCP (Fig. 1). For consistency, our live cell imaging and electron microscopy/tomography of mitochondria were limited to cortex cells in the root elongation zone. However, loss of membrane potential and mitochondrial recycling were observed in all cell types. To differentiate depolarized mitochondria, we pre-stained root cells with tetramethylrhodamine ethyl ester (TMRE), a fluorescent dye sensitive to membrane potential^15^. Normal mitochondria are seen as yellow puncta from dual fluorescence emitted from GFP and TMRE, while depolarized mitochondria will be green as TMRE will not fluoresce upon membrane depolarization.

**Figure 1.**
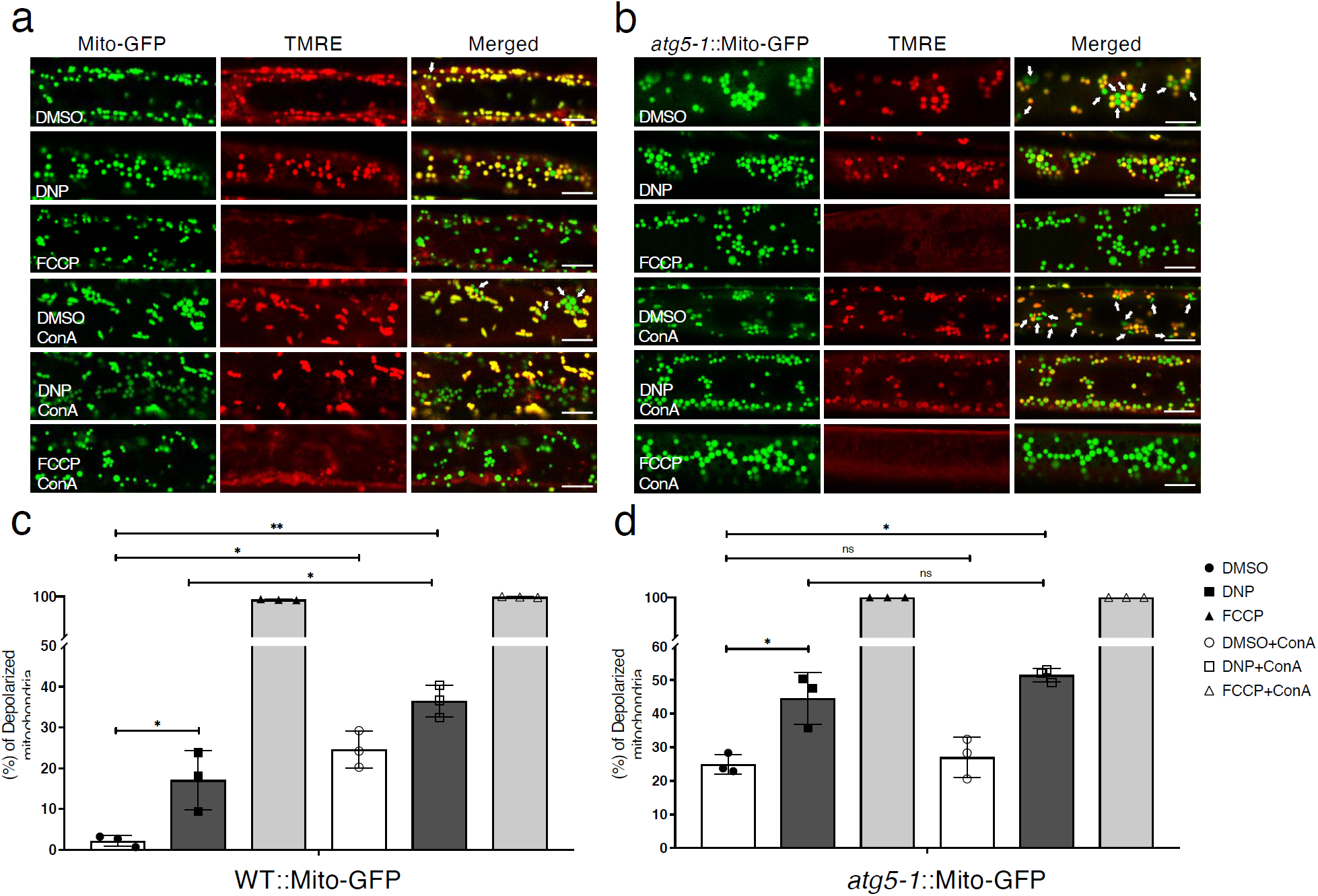
Arabidopsis root cells accumulate depolarized mitochondria upon uncoupler treatments. **a,b**, Uncoupler treatment induced mitochondria depolarization. Confocal micrographs of wild-type **(a)** and *atg5-1* **(b)** root cells expressing a mitochondrion-targeted GFP (Mito-GFP) after uncoupler treatment. Mitochondria were prestained with TMRE. Normal mitochondria exhibit yellow fluorescence while depolarized mitochondria exhibit green fluorescence (arrows) in DMSO or DMSO + ConA panels in the merged image columns. Note that most mitochondria are round. Scale bars, 8 μm. **c,d**, Histograms illustrating the percentage of depolarized mitochondria in wild type (WT) and *atg5-1* root cells expressing Mito-GFP at each treatment conditions. Bars represent the mean (± SD) of three biological replicates, each generated with three technical replicates. About 500 mitochondria from 10 cells (five root samples) were counted per condition. Asterisks (*) denote significant differences in depolarized mitochondria percentages relative to DMSO control group under each condition (unpaired t-test, *p<0.05, **p<0.01, ns, no significant difference).

Under normal conditions, depolarized mitochondria were rare in Mito-GFP roots (Fig. 1a). When Mito-GFP roots were incubated with DNP (50 μM) for 1 hr, numbers of depolarized mitochondria increased significantly (Fig. 1a,c). Inactivation of a core macroautophagy gene, *ATG5*, in the Mito-GFP line (*atg5-1∷*Mito-GFP) led to accumulation of more depolarized mitochondria in DNP-treated as well as untreated roots, indicating that removal of depolarized mitochondria requires *ATG5* (Fig. 1b,d). Addition of Concanamycin A (ConA), an inhibitor of vacuolar H^+^-ATPase that disrupts protein transport to the vacuole^16^, led to further build-up of mitochondria lacking membrane potential in Mito-GFP roots, indicating vacuole is the final destination for these depolarized mitochondria. Importantly, ConA did not lead to a similar build-up of depolarized mitochondria in *atg5-1*∷Mito-GFP roots (Fig. 1d, Extended Data Fig. 1). These observations agree with the inhibition of autophagy by ConA^17,18^ and suggest that depolarized mitochondria are recycled via the macroautophagy machinery in *Arabidopsis* root cells.

FCCP was a more potent uncoupler than DNP, depolarizing almost all mitochondria at a lower concentration after 1 hr (Fig. 1). In the following analyses, however, we employed DNP to trigger mitophagy, because its slower action facilitated the monitoring of the mitophagy dynamics via cell biological and biochemical assays.

### Uncoupler treatments induce autophagy

To examine whether autophagosome formation is induced following the uncoupler stress, we visualized a member of the *Arabidopsis* ATG8 family, ATG8e, fused with a YFP (YFP-ATG8e). ATG8 is widely used as a marker for autophagosomes in all eukaryotes^19,20^. Under normal conditions we rarely observed puncta; most of the ATG8 signal was diffuse. However, The YFP-ATG8e foci multiplied in root cells after 1 hr of DNP treatment, and their numbers continued to increase at later time points (Fig. 2a,b). Upon activation of autophagy, ATG8 becomes conjugated to phosphatidylethanolamine (PE) by a complex containing ATG5, and affixed to the limiting membrane of autophagosomes^17^. The lipidated form of ATG8 runs faster in western blots, and the ratio between lipidated and unlipidated ATG8 is used as a proxy to measure autophagy^16^. Immunoblot analyses with ATG8 antibody revealed only a faint upper band in untreated wild type (WT) cells (Fig. 2c). This band became more abundant by DNP treatment, especially in the membrane fraction. Samples incubated with phospholipase D (PLD) or samples from *atg5-1* mutants lacked the upper band (Fig. 2d). Altogether, these results suggest that ATG8 lipidation is induced upon DNP treatment in an *ATG5* dependent manner.

**Figure 2.**
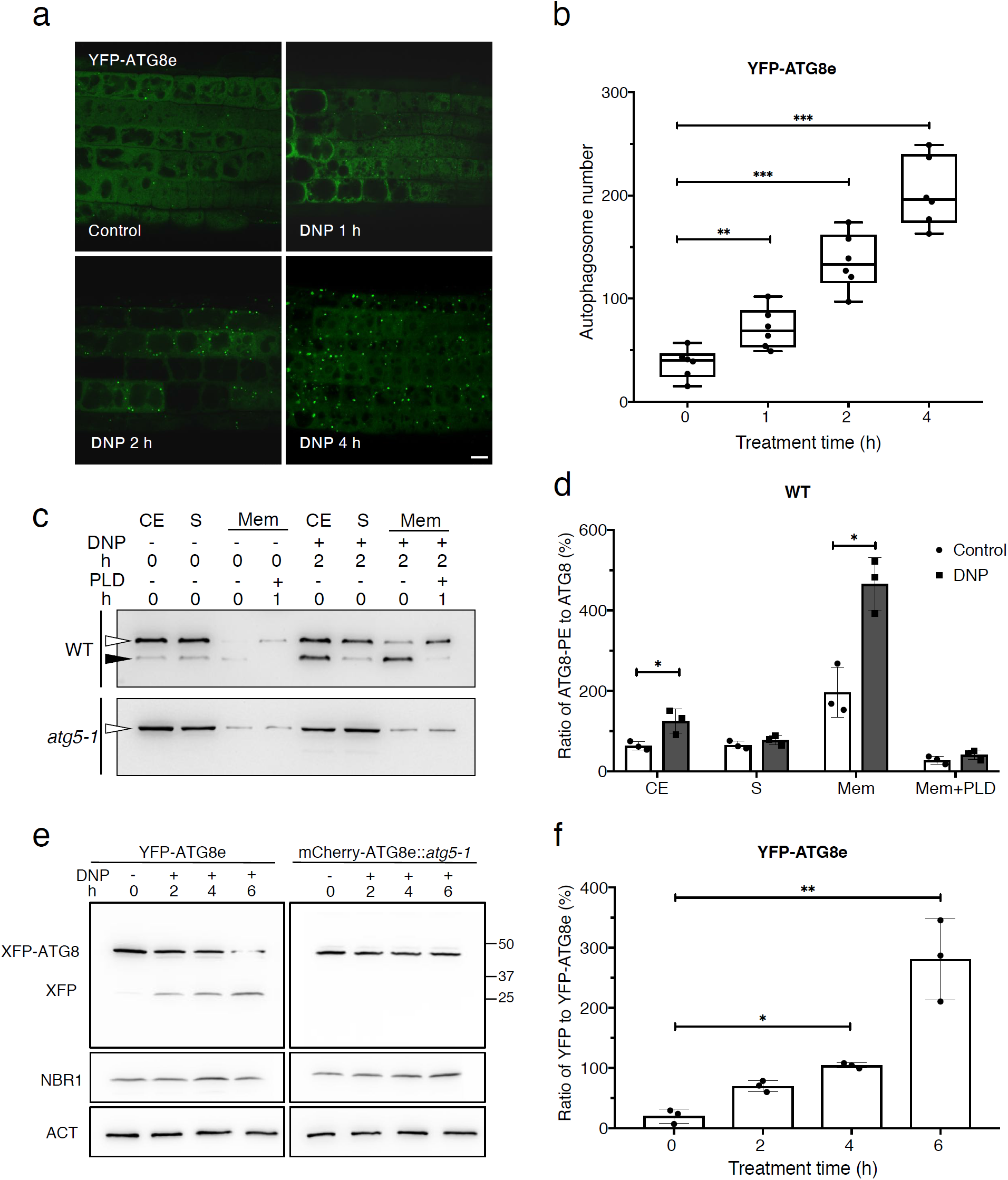
Uncoupler treatments induce autophagy. **a**, DNP treatment induces autophagosome formation. *Arabidopsis* YFP-ATG8e seedlings were incubated in DNP solution for varying periods (0-4 h) before imaging. **b**, Quantification of the number of autophagosomes from more than five independent root samples (unpaired t-test, **p<0.01, ***p<0.001). Scale bars, 8 μm. **c**, Uncoupler treatment activates ATG8 lipidation. Protein crude extracts (CE) were prepared from *Arabidopsis* root cells following incubation in DNP for 2 hours. Soluble (S) and membrane (Mem) fractions were separated and examined by immunoblot analysis with an anti-ATG8 antibody. White arrowheads mark ATG8. An additional polypeptide recognized by the antibody is enriched in the membrane fraction when cells are incubated with DNP (black arrowhead). **d**, Histograms illustrating polypeptide intensity ratios of lipidated ATG8 to free ATG8 in **(c).** Bars represent the mean (± SD) of three biological replicates. **e**, ATG8 cleavage assays of DNP treated WT and *atg5-1* seedlings expressing YFP-ATG8e or mCherry-ATG8e, respectively. Protein extracts were prepared from *Arabidopsis* seedlings exposed to DNP (50 μM) for the indicated time periods and subjected to immunoblot analysis with anti-GFP or anti-mCherry antibodies. NBR1 and Actin were used as control. **f**, Histograms illustrating the polypeptide intensity ratios of free YFP to YFP-ATG8e in **(d).** Bars represent the mean (± SD) of three biological replicates. (unpaired t-test, *p<0.05, **p<0.01).

To further confirm that the uncoupler stress induces autophagosome formation and vacuolar delivery, we performed GFP cleavage assay with the YFP-ATG8e line and *atg5-1* mutant line expressing an mCherry-ATG8e chimeric protein (mCherry-ATG8e∷*atg5-1*). A free YFP polypeptide was detected in YFP-ATG8e samples and its amount increased over time with a concomitant drop in YFP-ATG8e (Fig. 2e,f). No free mCherry was discerned in the immunoblot of mCherry-ATG8e::*atg5-1* by an anti-mCherry antibody (Fig. 2e). Excitingly, DNP treatment did not affect the autophagic flux of an aggrephagy receptor, Neighbor of BRCA1 (NBR1)^21^ (Fig. 2e). These data suggest that uncoupler treatment activates a selective autophagy pathway to recycle depolarized mitochondria.

### Damaged mitochondria are selectively engulfed by autophagosomes in uncoupler treated root cells

We then examined if DNP induced autophagosomes were indeed engulfing mitochondria by staining YFP-ATG8e root cells with MitoTracker Red (MTR). Under a confocal microscope, YFP-positive puncta were seen in the vicinity of mitochondria (Fig. 3a). In higher magnification micrographs, we were able to identify ATG8e-specific fluorescent structures resembling open pouches that contain mitochondria (arrowheads, Fig. 3b). We were also able to observe mitochondria that were entirely surrounded by YFP-ATG8e rings (white arrow, Fig. 3b). In time-lapse live cell imaging, YFP-ATG8e pockets partially enclosing a mitochondrion were seen to grow, eventually encapsulating the mitochondrion over 300 seconds (Fig. 3c). These autophagic compartments (i.e., mitophagosomes) were approximately 1-2 μm in diameters and each carried a single mitochondrion. MTR stains depolarized mitochondria better than TMRE but does not concentrate in mitochondria with no membrane potential^22^. In this vein, empty autophagosomes matching the size of mitophagosomes (grey arrow in Fig. 3b) in our micrographs could correspond to mitophagosomes carrying mitochondrial corpses.

**Figure 3.**
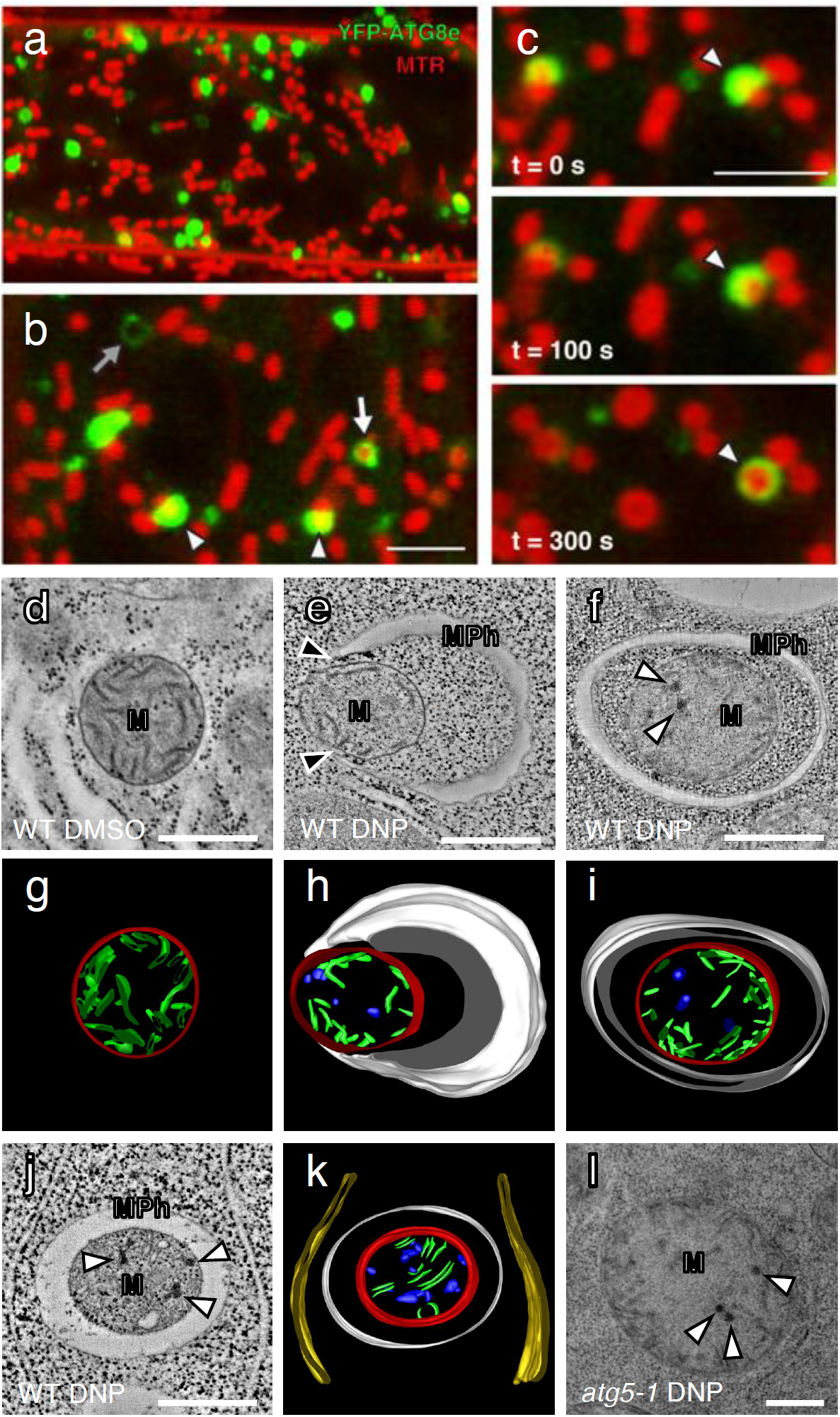
Depolarized mitochondria are selectively engulfed by autophagosomes in uncoupler-treated *Arabidopsis* root cells. **a,b**, DNP treatment induces mitophagy in *Arabidopsis* root cells. Confocal micrographs of *Arabidopsis* root cells expressing YFP-ATG8e stained with MitoTracker Red (MTR) and incubated with DNP for 1 hour. The mitochondria that associate with YFP-ATG8e are indicated with the arrowheads in panel (**b**). The mitochondria that are completely engulfed by ATG8e fluorescence is marked with a white arrow. Empty YFP fluorescence circles were also observed (grey arrow in panel **b**). Scale bars, 5 µm. **c**, Time lapse imaging of a mitophagy event in an *Arabidopsis* root tip cell treated with DNP. The mitochondrion is engulfed by YFP-ATG8e over 5 minutes. Scale bar, 5 µm. **d-f,j,l**, Transmission electron micrographs of mitochondria (M) in *Arabidopsis* WT or *atg5-1* root cells incubated with DMSO or DNP. Mitochondria phagophores (MPh) assemble in the vicinity of the mitochondria. Note that the phagophore tips (black arrow) are in contact with the mitochondrial surface in **(e). g-i,k**, Three-dimensional models of the mitophagosome (MPh) and its mitochondrial cargo (M) based on the tomogram in **(d-f,j,l)**. Mitophagosome (white), mitochondria outer membrane (red), mitochondria cristae (green), damaged cristae formed aggregates (blue) and ER (yellow) are modeled. White arrows indicate dark aggregates in the matrix of compromised mitochondria. Scale bars, 500 nm.

A time-lapse movie revealed that a small YFP-ATG8e puncta arose near MTR stained mitochondria and elongated to be a semicircle capturing a mitochondrion. This process took about 10 mins (Extended Data Fig. 2a). Another video documented an incomplete mitophagosome that expanded to enclose a mitochondrion fully (Extended Data Fig. 2b). Elongating tips of phagophores stayed in contact with the mitochondrial surface throughout their growth (Extended Data Fig. 3a). From our live cell microscopy data, we estimated that it takes about 15 min for mature mitophagosomes to develop from an initial YFP-ATG8 spot on a mitochondrion.

We then cryofixed root samples and performed transmission electron microscopy (TEM) analysis. Mitochondria in control samples had smooth cristae that are evenly dispersed in the matrix (Fig. 3d,g). By contrast, mitochondria in DNP treated cells had electron-dense precipitates in their matrix, some of which were engulfed by double membraned mitophagosomes (Fig. 3 e-k). Serial section TEM of mitophagosomes showed that one mitochondrion was contained per autophagosome, and no other organelles were identified in autophagosomes (Extended Data Fig. 3b,c), in agreement with the live cell imaging results (Fig. 3b,c). Electron tomography analysis revealed that mitochondria sequestered in mitophagosomes have more dark precipitate but less cristae in the matrix than free mitochondria (Fig. 3g-k). *Arabidopsis atg5-1* root cells had many mitochondria exhibiting the signs of the internal precipitates, but they were not associated with mitophagosomes (Fig. 3l). Altogether these results show that plant cells selectively recycle damaged mitochondria via autophagy that involves ATG5.

To further investigate how mitochondria are degraded during uncoupler induced mitophagy, we used immunoblot assays to assess the levels of various mitochondrial proteins in *Arabidopsis* WT and *atg5-1* mutant lines. We treated *Arabidopsis* seedlings with DNP for one to four hours (D1-D4). To distinguish between uncoupler induced mitophagy and bulk autophagy, we used nitrogen starvation as control (Extended Data Fig. 4). In contrast to the endoplasmic reticulum protein Cycloartenol-C24-methyl transferase (SMT1), levels of outer mitochondrial membrane (OMM) proteins peripheral-type benzodiazepine receptor (PBR) and voltage dependent anion channel 1 (VDAC1); inner mitochondria membrane (IMM) proteins cytochrome oxidase subunit II (COXII) and L-galactono-1,4-lactone dehydrogenase (GLDH), and mitochondria matrix (MM) protein isocitrate dehydrogenase (IDH) were all reduced upon uncoupler treatment (Fig. 4a). The reduction in protein levels were due to vacuolar degradation as addition of ConA prevented the degradation (Fig. 4b). Furthermore, the degradation was dependent on ATG5, since protein levels did not change significantly in DNP treated *atg5-1* mutant specimens (Fig. 4a,c). Interestingly, OMM proteins showed higher levels of degradation in contrast to IMM and matrix proteins. Since, previous studies in mammalian cells showed OMMs could also be degraded via the proteasome^23^, we checked OMM protein levels of DNP treated cells following proteasome inhibition by MG132. Adding MG132 indeed stabilized only OMM proteins but not IMM or matrix proteins (Fig. 4d). Considered together with normalized quantification of protein levels (Fig. 4e,f), these experiments suggest upon loss of membrane potential, proteasome and autophagy cooperate to degrade OMM proteins, whereas IMM and matrix proteins are primarily degraded by autophagy.

**Figure 4.**
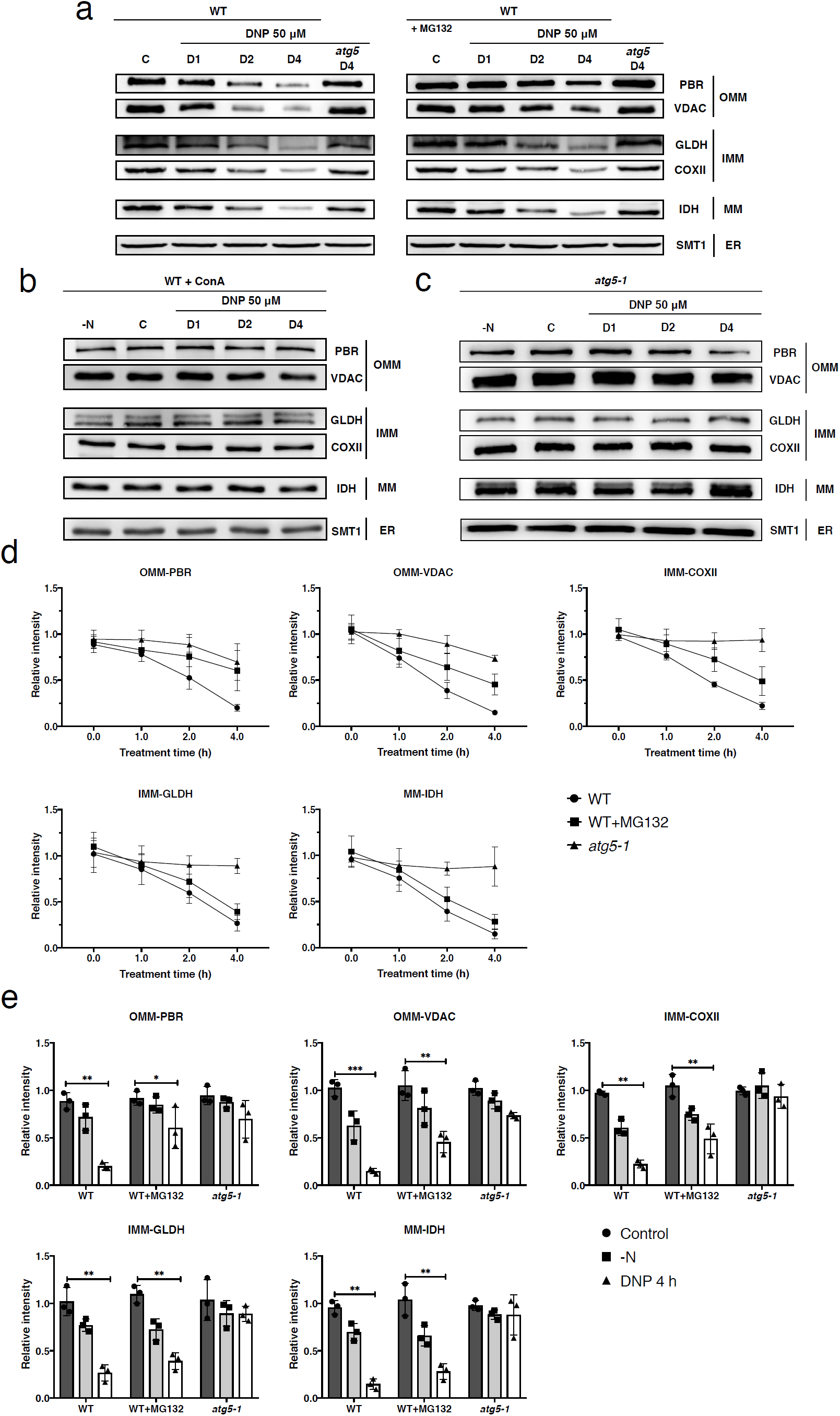
ATG5-dependent degradation of mitochondrial proteins in uncoupler-treated *Arabidopsis* root cells. **a-c**, Immunoblot blot analyses of uncoupler treatment induced mitochondrial protein degradation in *Arabidopsis* WT and *atg5-1* seedlings. WT and *atg5-1* mutant *Arabidopsis* roots were incubated in DNP solution for 1 to 4 hours (D1-D4) or nitrogen starvation (-N) solutions for 1 day. Mitochondrial outer membrane, mitochondrial matrix, and endomembrane fractions were isolated and subjected to immunoblot analyses. For outer mitochondrial membrane (OMM) proteins, PBR and VDAC1, for inner mitochondrial membrane (IMM) COXII and GLDH, and for mitochondrial matrix IDH were analysed. Concanamycin A (ConA) was added to the treatment solution to test for vacuolar function in mitochondrial protein degradation. Proteasome inhibitor MG132 was added to test the involvement of proteasomes in the recycling of mitochondrial membrane proteins **(c)**. Note that an ER protein, SMT1 was not affected by DNP treatment. Equal amounts of protein extracts were analysed in the immunoblots shown. **d**, Line charts illustrating degradation rates of OMM proteins (PBR and VDAC1), IMM proteins (COXII and GLDH), and MM protein (IDH) in WT treated with DNP (WT), *atg5-1* treated with DNP (*atg5-1*), and WT treated with DNP and MG132 (WT+MG132). **e**, Histograms illustrating the levels of mitochondria membrane proteins under the three treatment conditions, DMSO 4h (Control), DNP 4 h, and nitrogen starvation (-N) in WT and *atg5-1* root cells. The polypeptide intensity values were normalized to that of the loading control (SMT1). Bars represent the mean (± SD) and the asterisks (*) indicate decreases in polypeptide readouts significantly from that of the control (C) point (unpaired t-test, *P < 0.05, **P < 0.01, ***P < 0.01).

### *Friendly* is essential for uncoupler induced mitophagy

We then wanted to identify molecular players that mediate mitophagy in plants. Previous studies have shown that for both ubiquitin dependent and independent mitophagy pathways, mitochondrial network needs to be fragmented^24,25^. In *Arabidopsis*, Friendly (FMT), a clustered mitochondria (CLU) family protein, has been shown to play important roles in mitochondrial dynamics^26^. However, whether it plays a role in mitophagy wasn’t addressed. YFP-FMT exhibited a diffuse cytosolic pattern under normal conditions. Interestingly, upon DNP treatment, YFP-FMT localized to puncta that colocalized with mitochondria (Fig. 5a). Pull down experiments showed that uncoupler treatment led to specific association of YFP-FMT with ATG8, suggesting that YFP-FMT localizes to mitophagosomes (Fig. 5b). This prompted us to analyse mitophagy in *fmt* mutant. Consistent with our previous findings presented in Fig. 3, live cell imaging of GFP-tagged mitochondria in mCherry-ATG8e expressing WT cells showed ATG8e labelled vesicles engulfed mitochondria upon uncoupler treatment. However, in *fmt* mutant, although ATG8 puncta were associated with the mitochondria, we did not observe engulfment of mitochondria within autophagosomes (Fig. 5c). Further analyses of mitophagosomes in the *fmt* mutant using TEM revealed aberrant mitophagosomes with disconnected edges (Fig. 5c).

**Figure 5.**
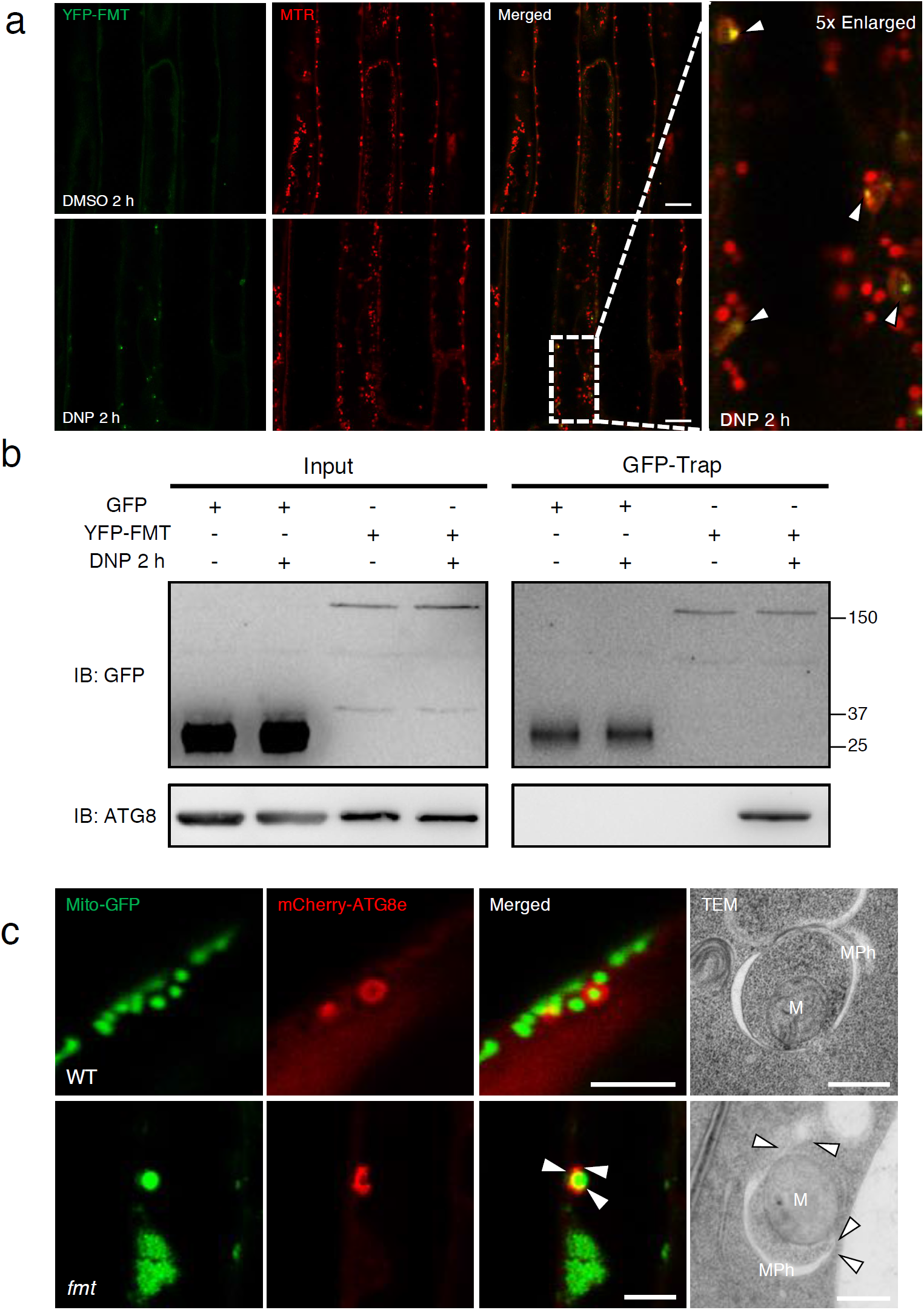
Friendly associates with damaged mitochondria and ATG8 upon uncoupler treatment. **a**, DNP treatment induces recruitment of Friendly (FMT) to damaged mitochondria. Confocal micrographs of *Arabidopsis* root cells expressing a YFP-tagged FMT (YFP-FMT) prestained with MTR. FMT-YFP seedlings were incubated with DMSO or DNP for 1 hour prior to imaging. Scale bars, 8 μm. **b**, FMT associates with ATG8 upon DNP treatment. *Arabidopsis* root cells expressing YFP-FMT were incubated with DMSO or DNP for 2 hours and then subjected to immunoprecipitation with GFP-trap followed by immunoblotting with indicated antibodies. **c**, Confocal micrographs of *Arabidopsis* WT and *fmt* root cells expressing mitochondria-targeted GFP (Mito-GFP) and mCherry-targeted ATG8e (mCherry-ATG8e). WT or *fmt* plants seedlings were incubated with DNP for 1 hour prior to imaging. Scale bars, 8 μm. Transmission electron microscopy (TEM) photos show mitochondrial phagophores (MPh) assemble in the vicinity of the mitochondria (M) under the condition for the experiment in **(a)**. Arrowhead point to defective phagophores in confocal and TEM micrographs. Scale bars, 500 nm.

To further test the role of FMT in mitophagy, we performed live cell imaging and immunoblot based autophagic flux experiments. Staining of depolarized mitochondria with TMRE upon uncoupler treatments revealed accumulation of damaged mitochondria in the cytosol of *fmt* mutants (Fig. 6a). Quantification of depolarized mitochondria showed *fmt* mutants accumulated significantly more mitochondria in contrast to WT cells (Fig. 6b). Furthermore, morphometric analyses of mitochondria in TEM micrographs from WT, *fmt* and *atg5-1* mutants showed that mitochondria were significantly larger in *fmt* and *atg5-1* mutants (Extended Data Fig. 5). Finally, immunoblot analyses using mitochondrial compartment specific antibodies also showed a delay of mitochondrial protein degradation in *fmt* mutant (Fig. 6c). Comparative analyses of the polypeptide intensities indicated that although *fmt* mutant had a significant defect in mitochondrial protein recycling (Fig. 6d), it was not as severe as the *atg5-1* mutant (Fig. 4d,e), suggesting, in the absence of FMT, compensatory pathways prevent accumulation of the toxic damaged mitochondria in the cell. Altogether these experiments suggest FMT is required for mitophagy in plants.

**Figure 6.**
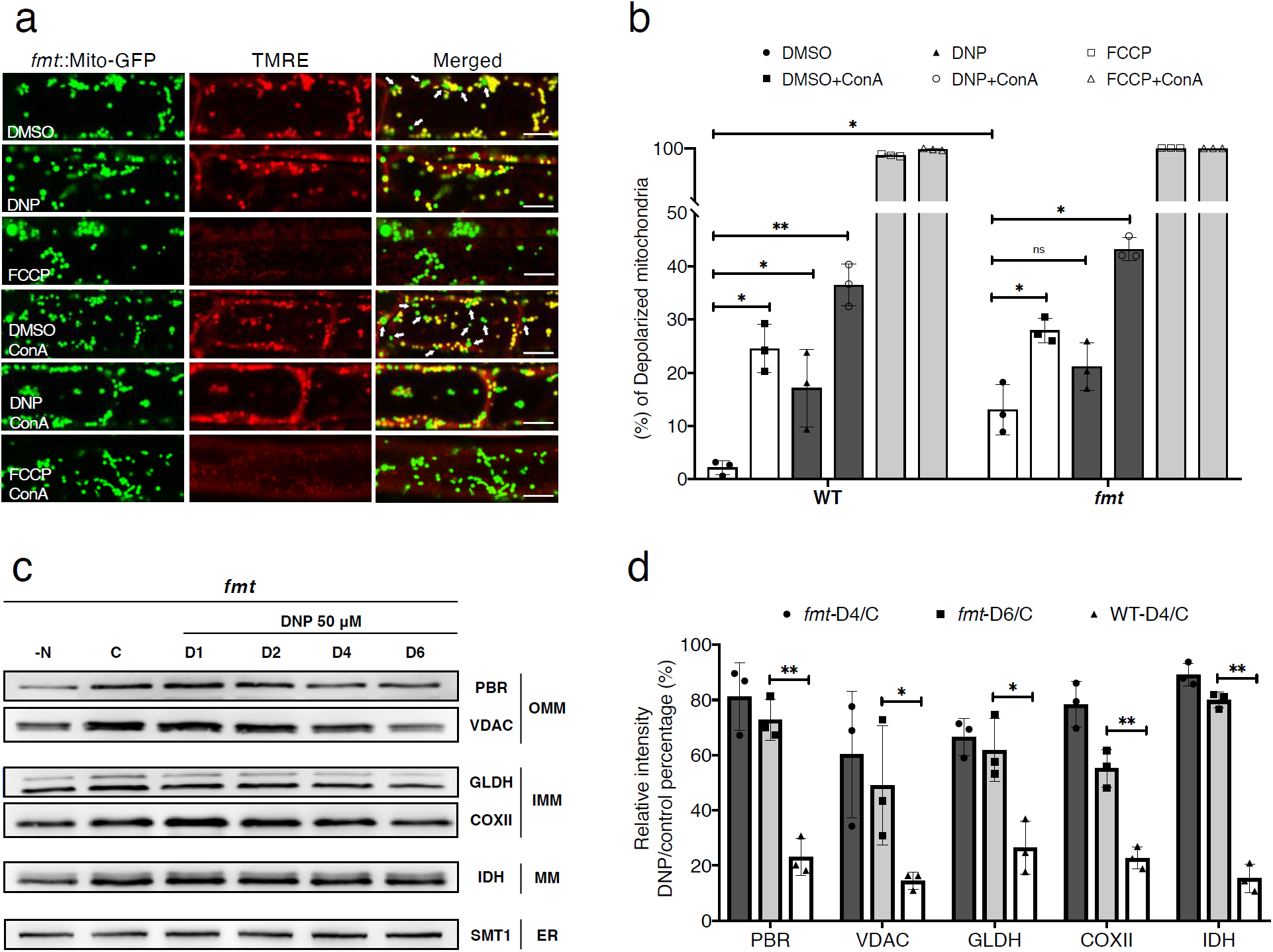
*Arabidopsis friendly* mutant have defects in clearance of depolarized mitochondria. **a**, Uncoupler treatment induced mitochondria depolarization in *Arabidopsis friendly* mutant (*fmt)*. Confocal micrographs of *fmt* root cells expressing a mitochondrion-targeted GFP (Mito-GFP), stained with TMRE. Normal mitochondria exhibit yellow fluorescence and depolarized mitochondria exhibit green florescence, denoted with arrows in DMSO or DMSO +ConA panels in the merged image column. *Arabidopsis* seedings were incubated with DMSO, DNP or FCCP with or without ConA for 1 hour prior to imaging. Scale bars, 8 μm. **b**, Histograms illustrating the percentage of depolarized mitochondria for each treatment conditions. Bars represent the mean (± SD) of three biological replicates. About 500 mitochondria from 10 cells (five root samples) were counted per condition. An asterisk (*) represents a significant difference of depolarized mitochondria percentage in each treatment relative to DMSO control group (unpaired t-test, *p<0.05, **p<0.01). **c**, Uncoupler treatment induced mitochondrial protein degradation in *fmt* mutant. *Arabidopsis fmt* mutant roots were incubated in DNP solutions for 1 hour to 6 hours (D1-D6) or nitrogen starvation (N-) solutions for 1 day. For outer mitochondrial membrane (OMM) proteins, PBR and VDAC1, for inner mitochondrial membrane (IMM) COXII and GLDH, and for mitochondrial matrix IDH were examined with immunoblot analysis. Note that an ER protein, SMT1, was not affected by DNP treatment. **d**, Histograms illustrating the levels of mitochondrial membrane protein degradation for the DNP treatment (DNP 4 h, DNP 6 h) in WT and *fmt* root cells. The polypeptide intensity values were normalized with that of the loading control (SMT1) and the percentage of DNP treatment group to control group are quantitated. Bars represent the mean (± SD) and the asterisks (*) indicate the significantly difference in polypeptide readouts between *fmt* and WT groups (unpaired t-test, *P < 0.05, **P < 0.01).

### Cotyledon greening during de-etiolation is affected in *atg5-1* mutant seedlings

In germinating seeds, under darkness, proplastids transform into etioplasts that are characterized by paracrystalline arrays of prolamellar bodies in their stroma^27^. The etioplast quickly transforms into the chloroplast upon exposure to light, a major developmental transition termed de-etiolation. Since mitochondrial and chloroplast functions are tightly interconnected, we hypothesized that mitochondrial population may also undergo remodelling during de-etiolation. First, we monitored greening of cotyledons when dark grown seedlings were exposed to light. Green pigment levels increased gradually over a 12 hr period, with a significant rise at eight hours after illumination (Fig. 7a,b). We then measured amounts of mitochondrial proteins in greening cotyledon cells and showed that their levels also drop at 8 hr time point, suggesting an accelerated removal of mitochondria (Fig 7c). Consistently, mitophagosomes were most frequently detected in cotyledon cell sections from 6 and 8 hr samples under TEM (Fig. 7d). Cotyledon greening was severely affected in *atg5-1* mutant seedlings and no mitophagosomes were discerned in their cotyledon cells (Fig. 7 a,b,e). These results indicate de-etiolation is a physiological stress condition for mitochondria where mitophagy is induced to mediate mitochondrial turnover that underlies light activated cotyledon development.

**Figure 7.**
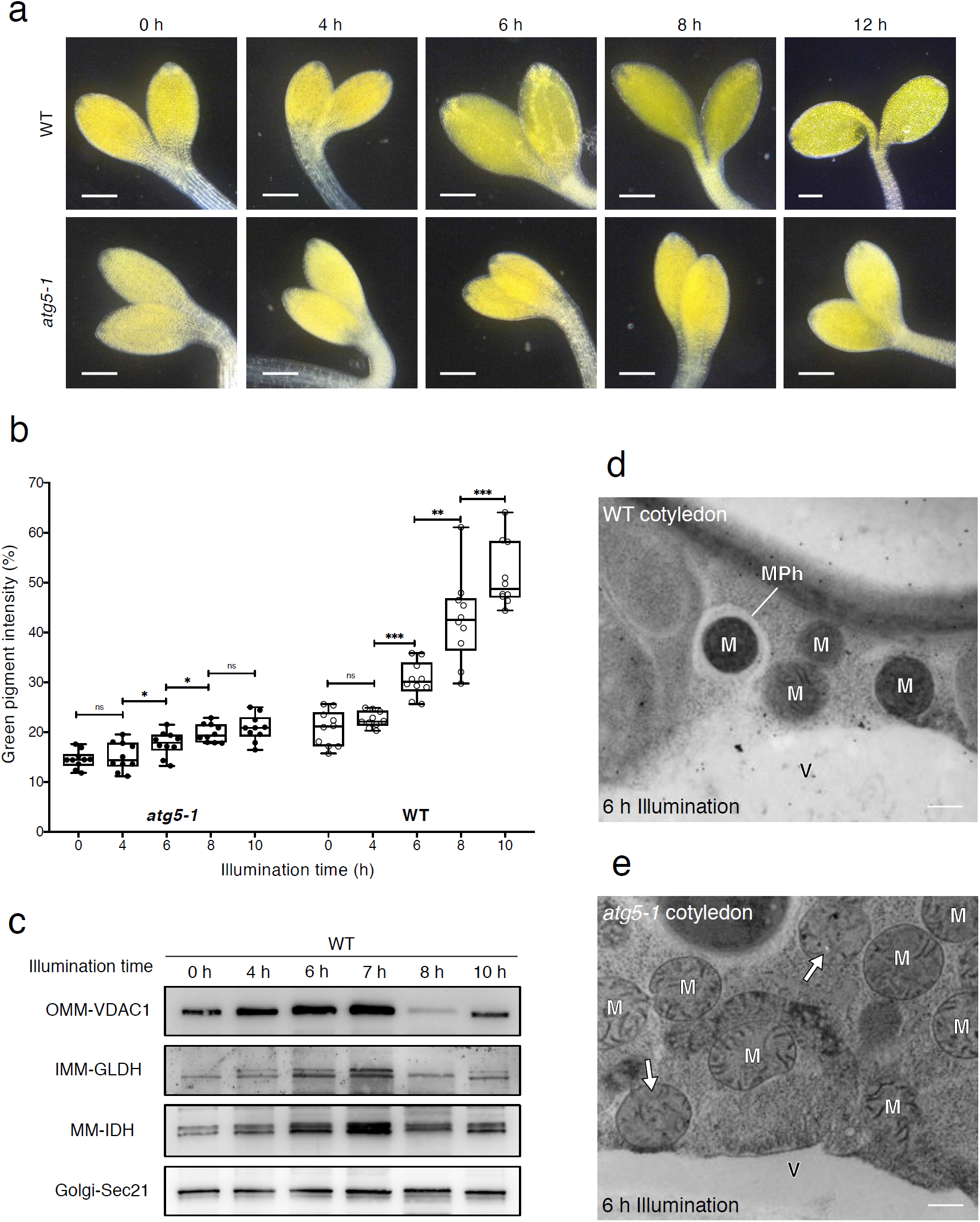
Mitophagy is triggered during de-etiolation of *Arabidopsis* seedlings. a, Cotyledon greening after light exposure in WT and atg5-1 Arabidopsis. Darkfield stereo microscopy photos showing Arabidopsis WT and atg5-1 mutant cotyledon at multiple time points (0-12 h) after illumination. Arabidopsis seedlings were grown darkness before the experiment. Scale bars, 1 mm. b, Quantification of the green pigments in cotyledons after light exposure. The green colour of ten Arabidopsis cotyledons for each time points were calculated from their photos and normalized against the dark background. (unpaired t-test, *p<0.05, **p<0.01, ***p<0.001, ns, no significant difference). c, Immunoblot analyses of mitochondrial proteins in Arabidopsis seedlings during de-etiolation. For outer mitochondrial membrane (OMM) proteins, PBR and VDAC1, for inner mitochondrial membrane (IMM) COXII and GLDH, and for mitochondrial matrix IDH were analysed as representative proteins. An Arabidopsis Golgi protein, coatomer subunit gamma (Sec21), was employed as the loading control. d,e, TEM images of a cluster of mitochondria (M) near the vacuole (V) in Arabidopsis WT and atg5-1 cotyledon cell after 6 hours of illumination. Mitochondria were seen to be surrounded by mitophagosome (MPh) in WT. By contrast, compromised mitochondria with dark aggregates were abundant in atg5-1 (arrows in e) but no mitophagosomes were associated with them. Scale bars, 500 nm.

## Discussion

Research in the last decade has transformed autophagy from a bulk degradation system to a highly selective cellular quality control pathway that rapidly removes toxic or superfluous macromolecules^28,29^. Especially organelles that get damaged due to metabolic and physiological stress conditions are mainly recycled via distinct selective autophagy pathways^6^. Consistently, in all the eukaryotes tested so far, autophagy is essential for adapting to environmental changes^12,30^. However, despite significant advances made in metazoan selective autophagy field, how plants recycle their organelles are still mostly unknown. Although the core autophagy machinery that mediates formation of the autophagosome is highly conserved in plants, selective autophagy receptors and adaptors that are responsible for recognition and recruitment of damaged organelles to the autophagosomes are not well conserved^12^. For example, there are up to eight different receptor proteins and dozens of accessory proteins that have been shown to mediate various mitophagy pathways in mammalian cells^3^. Homologs of most of those proteins are lacking in plant genomes, implying plant mitophagy have followed a different evolutionary path, likely due to the presence of another endosymbiont in the cell.

Here, we have established a detailed toolbox to study mitophagy in plants. We have shown that protonophore uncouplers specifically induce mitophagy similar to metazoans. Our findings also revealed high levels of mitophagy during de-etiolation, a fundamental developmental step that allows plants to survive day light after germination. Excitingly, our studies also revealed a molecular player, the Friendly protein, that is essential for mitophagy.

### Mitochondrial membrane potential is closely monitored by mitochondrial quality control pathways

As the primary producer of ATP in eukaryotic cells, healthy mitochondria must maintain the electrochemical potential. It is conceivable that the loss/reduction in the potential serves as a mark for the mitophagy machinery to recognize malfunctioning mitochondria. For example, in the PINK1/Parkin-dependent mitophagy pathway of mammalian cells, a mitochondrion-localized protein kinase, Pink1, is stabilized when the membrane potential drops and this leads to the activation of downstream effectors^31^. In *C. elegans* sperm mitochondria are rapidly removed from the oocyte via mitophagy upon fertilization^32,33^. Although the underlying mechanism of depolarization is still unknown, the paternal mitochondria that is recycled by mitophagy loses its membrane potential prior to mitophagy^15^. Our uncoupler treatments clearly demonstrated that membrane potential serves as a proxy for mitochondrial health across different organisms and loss of membrane potential triggers mitophagy. However, how plants tag depolarized mitochondria for mitophagy needs to be further investigated.

Mitochondria in mammalian and yeast cells undergo cycles of fusion and fission. This mitochondrial dynamics collaborate with mitophagy; fusion can rescue damaged mitochondria by diluting their injuries while fission singles out aberrant mitochondria for mitophagy. It was shown that mitochondria fusion is inhibited when the mitochondrial membrane potential is dissipated by uncouplers^34^. Mitochondria in Arabidopsis root cells are mostly round, indicating that mitochondria fission dominates over fusion^35^. Therefore, our system is not suited for investigating the link between mitochondria’s membrane potential and their fusion/fission dynamics. Some plant cells including seed cells after germination or shoot apical meristem cells have elongated mitochondria constituting a network^36,37^. These cells are better for testing whether mitochondria fragment in response to uncoupler stresses and studying roles of mitochondria fission in plant mitophagy.

### Ultrastructural features of compromised mitochondria and mitophagosomes in *Arabidopsis* root cells

Mitochondria with dark aggregates and shrivelled cristae in the matrix appeared after incubation with uncouplers. They were also abundant in de-etiolating cotyledon cells and fmt mutant cells (Fig. 3 and Extended Data Fig. 1). These impaired mitochondria were rare in DMSO control samples but frequently observed in *atg5-1* mutant samples in TEM images. Considering that they are specifically targeted by and enclosed in mitophagosomes, they correspond to depolarized mitochondria being recycled. Sperm mitochondria in cryofixed *C. elegans* oocytes also exhibited similar ultrastructural features^15,38^. However, we did not observe large ruptures in mitochondrial membranes reported in TEM analysis of mammalian mitophagy where cells were preserved by chemical fixation^39^.

An intriguing feature that we noticed in our TEM analysis is that tips of the elongating phagophores were in contact with the mitochondrial surface (Fig. 3e, arrowheads), This observation is consistent with the YFP-ATG8e fluorescence that expanded tightly over mitochondria in the time-lapse recordings (Fig. 3c). The affinity between the autophagosome membrane and mitochondria explains why mitophagosomes usually have one damaged mitochondrion, and they do not capture other organelles. It also suggests specific protein-protein interactions regulate the two membranes, which need to be investigated further.

### FMT is essential for mitophagy in plants

FMT is required for normal mitochondria distribution in the plant cell, possibly regulating association between individual mitochondria before fusion^26^. We have shown that DNP-induced mitochondria recycling is affected in *fmt* mutant cells, and FMT associates with ATG8 upon mitochondrial damage. These data suggest that FMT associates with autophagosomes during mitophagy. Because FMT is a protein shuttling between the cytosol and mitochondria, it is tempting to speculate that FMT may participate in an autophagy receptor or adaptor complex through which the core autophagy machinery is recruited to mitochondria. It is also possible that FMT plays a role in the repair of damaged mitochondria by controlling their interaction with the healthy mitochondria, and excessive accumulation of FMT triggers the onset of mitophagy.

FMT is a highly conserved protein with orthologs in evolutionarily distant eukaryotes including yeast and metazoans^40^. *fmt* mutants exhibit similar phenotypes where mitochondria form large clusters next to nucleus. The Drosophila FMT homolog Clueless positively regulates PINK1/Parkin dependent mitophagy by suppressing mitochondrial fusion^41^. Recent studies have shown that mammalian FMT homolog CLUH could bind RNA to form granules. CLUH granules regulate translation of mRNAs linked to metabolic activity and regulate mitophagy and metabolic reprogramming^42,43^. Plants lack PINK1/Parkin homologs, so whether FMT regulates mitophagy by forming stress activated granules or via a PINK/Parkin-like pathway need to be investigated further. Identification of FMT interacting proteins and RNA during nutrient starvation or uncoupler treatments could help us understand the role of FMT in mitophagy and mitochondrial quality control.

### Mitochondrial recycling mediate organellar reprogramming during de-etiolation

Reprograming the mitochondrial functions in response to nutrient availability is critical for cell survival^44^. In plant cells, photosynthesis in chloroplasts and respiration in mitochondria are coordinated for homeostasis of cellular energy levels and redox status^45,46^. Our results from greening Arabidopsis cotyledons indicated that de-etiolation involves a wave of mitochondrial turnover, probably for rewiring mitochondrial metabolic network for adapting to light condition and salvaging raw materials for chloroplast biogenesis.

In skotomorphogenic seedlings, nutrients reserved in the seed are mobilized to sustain growth, and mitochondria are required for the anabolic processes. When light is available, the seedlings become photosynthetically active and capable of autotrophic growth. It was shown that the electron transport chain in mitochondria is slowed down in de-etiolating wheat leaves and chloroplasts and mitochondria are functionally more intertwined^47^. Accumulation of mitophagosomes and rapid changes in mitochondrial protein levels in the greening cotyledon suggest a cannibalization of pre-existing mitochondria. We speculate that the developmentally programmed mitophagy could facilitate modulation of the mitochondrial pool and biosynthesis of macromolecules for constructing new organelles. The lack of mitophagosomes and the delay in greening of the *atg5-1* mutant cotyledon agrees with the notion. Altogether, our findings on greening cotyledons present a clear example of inter-organelle communication, and how mitochondria-chloroplast crosstalk could underlie a major developmental transition in plants.

## Methods

### Materials and plant growth conditions

All the chemicals were purchased from Sigma-Aldrich (http://www.sigmaaldrich.com) or Thermo-Fisher (https://www.thermofisher.com) unless specified. YFP-ATG8e, mCherry-ATG8e, *atg5-1*, Mito-GFP, Mito-GFP::*fmt* and FMT-YFP seeds were described previously^26,48,49^. mCherry-ATG8e::*atg5-1*, Mito-GFP x mCherry-ATG8e and Mito-GFP x mCherry-ATG8e::*fmt* was obtained by crossing the previously established lines^26,50^. All transgenic lines were genotyped by PCR and homozygous lines were isolated before experiments. All *Arabidopsis* seeds were surface-sterilized and geminated on ½ Murashige and Skoog (MS) agar plate in a growth chamber at 21 °C with 16 h light–8 h dark except for de-etiolation experiments where seedlings were geminated under darkness.

### Protein extraction and immunoblot analysis

Isolation of mitochondrial membrane proteins and GFP cleavage assay were carried out as described previously^48,51^. Briefly, seven-day-old *Arabidopsis* seedlings were incubated in DMSO, 50 μM DNP, 50 μm MG132 or 1 μm ConA with indicated times in liquid half MS medium when necessary. Seedlings for the nitrogen starvation experiments were germinated on half MS medium agar plate and then transfer to liquid half MS medium without nitrogen for 1 day. All the protein samples were subjected to 15% SDS-PAGE. Primary and secondary antibodies were diluted in 1x phosphate buffered saline (PBS). Antibodies against GFP (Abcam), YFP (Agrisera), mCherry (Abcam), ATG8 (Agrisera), voltage-dependent anion channel 1 (VDAC1; Agrisera), peripheral-type benzodiazepine receptor (PBR; PhytoAB), cytochrome oxidase subunit II (COXII; Agrisera), L-galactono-1,4-lactone dehydrogenase (GLDH; Agrisera, PhytoAB) and isocitrate dehydrogenase (IDH; Agrisera), Cycloartenol-C24-methyl transferase (SMT1; Agrisera), Coatomer subunit gamma (Sec21p; Agrisera) were obtained from the indicated sources. We performed Student’s t test (one-tailed and unpaired test) with the triple replicate immunoblot data and the quantification of band intensities was performed using ImageJ (National Institutes of Health) and Microsoft Excel 2016, the graphs were made by Prism8 (GraphPad Software). Representative of at least three independant immunoblot results were shown in the figures.

### ATG8 delipidation assay

Protein extraction methods for ATG8-delipidation were described previously^52^. Seven-day-old *Arabidopsis* seedlings were incubated in 50 μM DNP for 2 hours before protein extraction. The total plant lysates were extracted in lysis buffer [50 mM Tris-HCl (pH 8.0), 150 mM NaCl, 0.5 mM Ethylenediaminetetraacetic acid (EDTA) and 1x Complete Protease Inhibitor Cocktail] and then centrifuged at 14 000 rpm for 10 min at 4 °C. The supernatant was centrifuged at 100 000 g for 1 h, with the membrane pellet then solubilized in lysis buffer containing 0.5% (v/v) Triton X-100. The solubilized membrane samples were incubated at 37°C for 1 h with 250 unit/ml of phospholipase D (PLD) or an equal volume of its buffer. Protein samples were subjected to 15% SDS-PAGE in the presence of 6 M urea and analyzed by immunoblot with anti-ATG8 antibody.

### Immunoprecipitation

Protein extraction and immunoprecipitation were performed as described previously^48^. Seven-day-old *Arabidopsis* seedlings were incubated in 50 μM DNP for 2 hours before protein extraction. Total plant lysates were centrifuged at 14000 rpm for 10 min at 4 °C. The supernatant was prepared in lysis buffer (10 mM Tris/HCl at pH 7.4, 150 mM NaCl, 0.5 mM EDTA, 5% glycerol, 0.2% Nonidet P-40, and 2 mM dithiobis [succinimidyl propionate] containing 1x Complete Protease Inhibitor Cocktail) and then incubated with GFP-TRAP agarose beads (ChromoTek) for 2 hours at 4°C. The beads were washed five times (4°C) in wash buffer (10 mM Tris/HCl, pH 7.5, 150 mM NaCl, and 0.5 mM EDTA with 1x Complete Protease Inhibitor Cocktail) and then eluted by boiling in 2x SDS sample buffer. Samples were separated by SDS-PAGE and analyzed by immunoblot using indicated antibodies.

### Confocal microscopy and image processing

Confocal fluorescence images were acquired using the Leica SP8 laser scanning confocal system with a 63x water lens. Seven-day-old *Arabidopsis* seedlings were incubated in DMSO, 50 μM DNP, 10 μM FCCP or 0.5 μM ConA with indicated times in liquid half MS medium when necessary before imaging. Tetramethylrhodamine ethyl ester (TMRE) and MitoTracker Red (MTR) were used to stain *Arabidopsis* root cell mitochondria at 500 nm for 10 mins. A sequential acquisition was applied when observing fluorescent proteins. Images were processed with Photoshop CC (https://www.adobe.com) and performed Student’s t test (one-tailed and unpaired test) with Microsoft Excel 2016 (https://www.microsoft.com/). The graphs were prepared with Prism8 (https://www.graphpad.com).

### TEM analysis, electron tomography, and 3d modeling

For TEM samples preparation, high-pressure freezing, freeze substitution, resin embedding, and ultramicrotomy were performed as described previously^53,54^. In brief, Seven-day-old *Arabidopsis* seedlings were incubated in DMSO or 50 μM DNP for indicated times and then rapidly frozen with an HPM100 high-pressure freezer (Leica Microsystems). The samples were freeze-substituted at −80°C for 72 h, and excess OsO4 was removed by rinsing with precooled acetone. After being slowly warmed up to room temperature over 48 h, root samples were separated from planchettes and embedded in Embed-812 resin (Electron Microscopy Sciences). Thin sections (100 nm thick) prepared from sample blocks of each time point were examined with a Hitachi 7400 TEM (Hitachi-High Technologies) operated at 80 kV.

For dual-axis tomography analysis, semi-thick sections (250 nm) were collected on formvar-coated copper slot grids (Electron Microscopy Sciences) and stained with 2% uranyl acetate in 70% methanol followed by Reynold’s lead citrate as described previously^55^. Tilt series were collected from 60° to - 60° (1.5° intervals) with a 200-kV Tecnai F20 intermediate voltage electron microscopy (https://www.fei.com/). Tomograms were reconstructed as described^56^. To generate models of complicated thylakoid membranes, we used the autocontour command (bio3d.colorado.edu/imod/doc/3dmodHelp/autox.html) of the 3dmod software package as explained in Keith and Kang (2017)^57^.

## Acknowledgements

We appreciate Dr. Xiaohong Zhuang (Chinese University of Hong Kong) for the *atg5 and atg7* mutant lines, David Logan and David Macharel for kindly sharing Mito-GFP and Friendly related seeds. We also thank Samantha Krasnodebski for help with generation of Arabidopsis lines. This work was supported by grants from the Research Grants Council of Hong Kong (GRF14126116, GRF14121019, C4012-16E, C4002-17G, and AoE/ M-05/12) and Cooperative Research Program for Agriculture Science & Technology Development (Project No. 0109532019) Rural Development Administration, Republic of Korea to B.-H.K., and Austrian Academy of Sciences and Austrian Science Fund (FWF): P32355 to Y.D.

## Author contributions

J.M., Y.D., and B.-H.K. conceived and designed the experiments. J.M. performed the confocal microscopy and stereomicroscopy. J.M., Z.L., and P.W. carried out electron microscopy/tomography analysis. J.M. and J.F. prepared 3D tomographic models. J.M. and W.M. performed immunoblot and pull-down experiments. J.Z., Y.Z., and N.G. did other experiments. J.M., J.Z., Z.L., P.W., L.J., Y.D., and B.-H.K. analysed the data. J.M., Y.D., and B.-H.K. wrote the paper.

## References

1. Youle, R. J. Mitochondria—Striking a balance between host and endosymbiont. Science (80-.). 365, (2019).

2. Broda, M., Millar, A. H. & Van Aken, O. Mitophagy: A Mechanism for Plant Growth and Survival. Trends Plant Sci. 23, 434–450 (2018).

3. Pickles, S., Vigié, P. & Youle, R. J. Mitophagy and Quality Control Mechanisms in Mitochondrial Maintenance. Curr. Biol. 28, R170–R185 (2018).

4. Palikaras, K., Lionaki, E. & Tavernarakis, N. Mechanisms of mitophagy in cellular homeostasis, physiology and pathology. Nat. Cell Biol. 20, 1013–1022 (2018).

5. Dikic, I. Proteasomal and Autophagic Degradation Systems.

6. Anding, A. L. & Baehrecke, E. H. Cleaning House: Selective Autophagy of Organelles. Dev. Cell 41, 10–22 (2017).

7. Nguyen, T. N., Padman, B. S. & Lazarou, M. Deciphering the Molecular Signals of PINK1 / Parkin Mitophagy. xx, 1–12 (2016).

8. Montava-Garriga, L. & Ganley, I. G. Outstanding Questions in Mitophagy: What We Do and Do Not Know. J. Mol. Biol. (2019) doi:10.1016/j.jmb.2019.06.032.

9. Georgakopoulos, N. D., Wells, G. & Campanella, M. The pharmacological regulation of cellular mitophagy. Nat. Chem. Biol. 13, 136–146 (2017).

10. Wauer, T., Simicek, M., Schubert, A. & Komander, D. Mechanism of phospho-ubiquitin-induced PARKIN activation. Nature 524, 370–4 (2015).

11. Lazarou, M. et al. The ubiquitin kinase PINK1 recruits autophagy receptors to induce mitophagy. (2015) doi:10.1038/nature14893.

12. Stephani, M. & Dagdas, Y. Plant Selective Autophagy - Still an uncharted territory with a lot of hidden gems. J. Mol. Biol. (2019) doi:10.1016/j.jmb.2019.06.028.

13. Li, F., Chung, T. & Vierstra, R. D. AUTOPHAGY-RELATED11 plays a critical role in general autophagy- and senescence-induced mitophagy in Arabidopsis. Plant Cell 26, 788–807 (2014).

14. Ashrafi, G. & Schwarz, T. L. The pathways of mitophagy for quality control and clearance of mitochondria. Cell Death Differ. 20, 31–42 (2013).

15. Zhou, Q. et al. Mitochondrial endonuclease G mediates breakdown of paternal mitochondria upon fertilization. Science (80-.). 353, 394–399 (2016).

16. Klionsky, D. J. et al. Guidelines for the use and interpretation of assays for monitoring autophagy (3rd edition). Autophagy 12, 1–222 (2016).

17. Marshall, R. S. & Vierstra, R. D. Autophagy: The Master of Bulk and Selective Recycling. Annu. Rev. Plant Biol. 69, 173–208 (2018).

18. Thompson, A. R., Doelling, J. H., Suttangkakul, A. & Vierstra, R. D. Autophagic Nutrient Recycling in Arabidopsis Directed by the ATG8 and ATG12 Conjugation Pathways. Plant Physiol. 138, 2097–2110 (2005).

19. Hanamata, S. et al. In vivo imaging and quantitative monitoring of autophagic flux in tobacco BY-2 cells. 1–11 (2013).

20. Kellner, R., De la Concepcion, J. C., Maqbool, A., Kamoun, S. & Dagdas, Y. F. ATG8 Expansion: A Driver of Selective Autophagy Diversification? Trends Plant Sci. (2016) doi:10.1016/j.tplants.2016.11.015.

21. Svenning, S., Lamark, T., Krause, K. & Johansen, T. Plant NBR1 is a selective autophagy substrate and a functional hybrid of the mammalian autophagic adapters NBR1 and p62 / SQSTM1. 3, 993–1010 (2011).

22. Kholmukhamedov, A., Schwartz, J. M. & Lemasters, J. J. Isolated mitochondria infusion mitigates ischemia-reperfusion injury of the liver in rats: mitotracker probes and mitochondrial membrane potential. Shock 39, 543 (2013).

23. Yoshii, S. R., Kishi, C., Ishihara, N. & Mizushima, N. Parkin mediates proteasome-dependent protein degradation and rupture of the outer mitochondrial membrane. J. Biol. Chem. 286, 19630–19640 (2011).

24. Lieber, T., Jeedigunta, S. P., Palozzi, J. M., Lehmann, R. & Hurd, T. R. Mitochondrial fragmentation drives selective removal of deleterious mtDNA in the germline. Nature 570, 380–384 (2019).

25. Twig, G. & Shirihai, O. S. The interplay between mitochondrial dynamics and mitophagy. Antioxidants Redox Signal. 14, 1939–1951 (2011).

26. El Zawily, A. M. et al. FRIENDLY Regulates Mitochondrial Distribution, Fusion, and Quality Control in Arabidopsis. Plant Physiol. 166, 808–828 (2014).

27. Ka, K., Mai, K., Gao, P. & Kang, B. Electron Microscopy Views of Dimorphic Chloroplasts in C4 Plants. 11, 1–7 (2020).

28. Mizushima, N. A brief history of autophagy from cell biology to physiology and disease. Nat. Cell Biol. 20, 521–527 (2018).

29. Pohl, C. & Dikic, I. Cellular quality control by the ubiquitin-proteasome system and autophagy. Science (80-.). 366, 818–822 (2019).

30. Levine, B. & Kroemer, G. Biological Functions of Autophagy Genes: A Disease Perspective. Cell 176, 11–42 (2019).

31. Wade Harper, J., Ordureau, A. & Heo, J. M. Building and decoding ubiquitin hains for mitophagy. Nat. Rev. Mol. Cell Biol. 19, 93–108 (2018).

32. Sato, M. & Sato, K. Degradation of Paternal Mitochondria. Science (80-.). 37, 1141–1144 (2011).

33. Cummins, J. M. et al. Postfertilization Autophagy of Sperm. 1, 1144–1148 (2011).

34. Malka, F. et al. Separate fusion of outer and inner mitochondrial membranes. EMBO Rep. 6, 853–859 (2005).

35. Arimura, S. Fission and fusion of plant mitochondria, and genome maintenance. Plant Physiol. pp. 01025.2017 (2017) doi:10.1104/pp.17.01025.

36. Sheahan, M. B., McCurdy, D. W. & Rose, R. J. Mitochondria as a connected population: Ensuring continuity of the mitochondrial genome during plant cell dedifferentiation through massive mitochondrial fusion. Plant J. 44, 744–755 (2005).

37. Seguí-Simarro, J. M. & Staehelin, L. A. Cell cycle-dependent changes in Golgi stacks, vacuoles, clathrin-coated vesicles and multivesicular bodies in meristematic cells of Arabidopsis thaliana: A quantitative and spatial analysis. Planta 223, 223–236 (2006).

38. Wang, Y. et al. Kinetics and specificity of paternal mitochondrial elimination in Caenorhabditis elegans. Nat. Commun. 7, 1–15 (2016).

39. Wei, Y., Chiang, W. C., Sumpter, R., Mishra, P. & Levine, B. Prohibitin 2 Is an Inner Mitochondrial Membrane Mitophagy Receptor. Cell 168, 224-238.e10 (2017).

40. Cox, R. T. & Spradling, A. C. Clueless, a conserved Drosophila gene required for mitochondrial subcellular localization, interacts genetically with parkin. DMM Dis. Model. Mech. 2, 490–499 (2009).

41. Wang, Z. H., Clark, C. & Geisbrecht, E. R. Drosophila clueless is involved in Parkin-dependent mitophagy by promoting VCP-mediated Marf degradation. Hum. Mol. Genet. 25, 1946–1964 (2016).

42. Pla-Martín, D. et al. CLUH granules coordinate translation of mitochondrial proteins with mTORC1 signaling and mitophagy. EMBO J. 39, 1–23 (2020).

43. Sheard, K. M., Thibault-Sennett, S. A., Sen, A., Shewmaker, F. & Cox, R. T. Clueless forms dynamic, insulin-responsive bliss particles sensitive to stress. Dev. Biol. 459, 149–160 (2020).

44. Spinelli, J. B. & Haigis, M. C. The multifaceted contributions of mitochondria to cellular metabolism. Nat. Cell Biol. 20, 745–754 (2018).

45. Raghavendra, A. S., Padmasree, K. & Saradadevi, K. Interdependence of photosynthesis and respiration in plant cells: interactions between chloroplasts and mitochondria. Plant Sci. 97, 1–14 (1994).

46. Van Lis, R. & Atteia, A. Control of mitochondrial function via photosynthetic redox signals. Photosynth. Res. 79, 133–148 (2004).

47. Garmash, E. V. et al. Expression profiles of genes for mitochondrial respiratory energy-dissipating systems and antioxidant enzymes in wheat leaves during deetiolation. J. Plant Physiol. 215, 110–121 (2017).

48. Zhuang, X. et al. A BAR-Domain Protein SH3P2, Which Binds to Phosphatidylinositol 3-Phosphate and ATG8, Regulates Autophagosome Formation in Arabidopsis. 1–21 (2013) doi:10.1105/tpc.113.118307.

49. Paszkiewicz, G., Gualberto, J. M., Benamar, A., Macherel, D. & Logan, D. C. Arabidopsis seed mitochondria are bioenergetically active immediately upon imbibition and specialize via biogenesis in preparation for autotrophic growth. Plant Cell 29, 109–128 (2017).

50. Zhuang, X. et al. ATG9 regulates autophagosome progression from the endoplasmic reticulum in *Arabidopsis*; Proc. Natl. Acad. Sci. 114, E426 LP–E435 (2017).

51. Marshall, R. S., Li, F., Gemperline, D. C., Book, A. J. & Vierstra, R. D. Autophagic Degradation of the 26S Proteasome Is Mediated by the Dual ATG8/Ubiquitin Receptor RPN10 in Arabidopsis. Mol. Cell 58, 1053–1066 (2015).

52. Chung, T., Phillips, A. R. & Vierstra, R. D. ATG8 lipidation and ATG8-mediated autophagy in Arabidopsis require ATG12 expressed from the differentially controlled ATG12A and ATG12B loci. Plant J. 62, 483–493 (2010).

53. Kang, B. H. Electron microscopy and high-pressure freezing of arabidopsis. Methods in Cell Biology vol. 96 (Elsevier Inc., 2010).

54. Wang, P., Chen, X., Goldbeck, C., Chung, E. & Kang, B. H. A distinct class of vesicles derived from the trans-Golgi mediates secretion of xylogalacturonan in the root border cell. Plant J. 92, 596–610 (2017).

55. Liang, Z. et al. Thylakoid-bound polysomes and a dynamin-related protein, FZL, mediate critical stages of the linear chloroplast biogenesis program in greening arabidopsis cotyledons. Plant Cell 30, 1476–1495 (2018).

56. Toyooka, K. & Kang, B.-H. Reconstructing Plant Cells in 3D by Serial Section Electron Tomography BT - Plant Cell Morphogenesis: Methods and Protocols. in (eds. Žárský, V. & Cvrčková, F.) 159–170 (Humana Press, 2014). doi:10.1007/978-1-62703-643-6_13.

57. Mai, K.K.K. & Kang, B.-H. Semiautomatic Segmentation of Plant Golgi Stacks in Electron Tomograms Using 3dmod. - Plant Protein Secretion: Methods and Protocols. in (eds. Liwen Jiang) 97–104 (Springer Science+Business Media, 2017). doi:10.1007/978-1-4939-7262-3_8

