## Supplemental Figures for "Friendly regulates membrane depolarization induced mitophagy in Arabidopsis"

### Slide 1
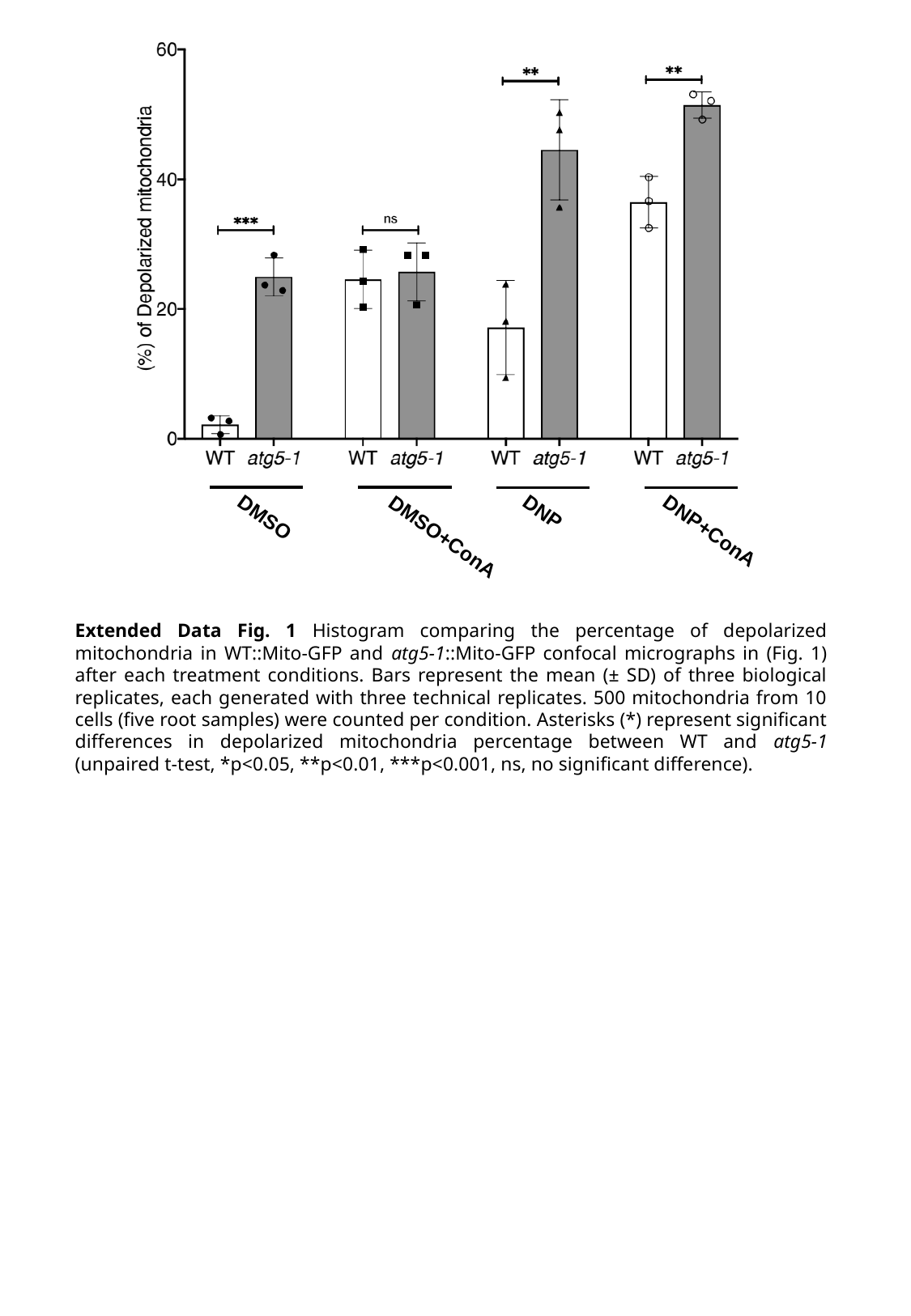

DNP
DMSO
DNP+ConA
DMSO+ConA
Extended Data Fig. 1 Histogram comparing the percentage of depolarized mitochondria in WT::Mito-GFP and atg5-1::Mito-GFP confocal micrographs in (Fig. 1) after each treatment conditions. Bars represent the mean (± SD) of three biological replicates, each generated with three technical replicates. 500 mitochondria from 10 cells (five root samples) were counted per condition. Asterisks (*) represent significant differences in depolarized mitochondria percentage between WT and atg5-1 (unpaired t-test, *p<0.05, **p<0.01, ***p<0.001, ns, no significant difference).

### Slide 2
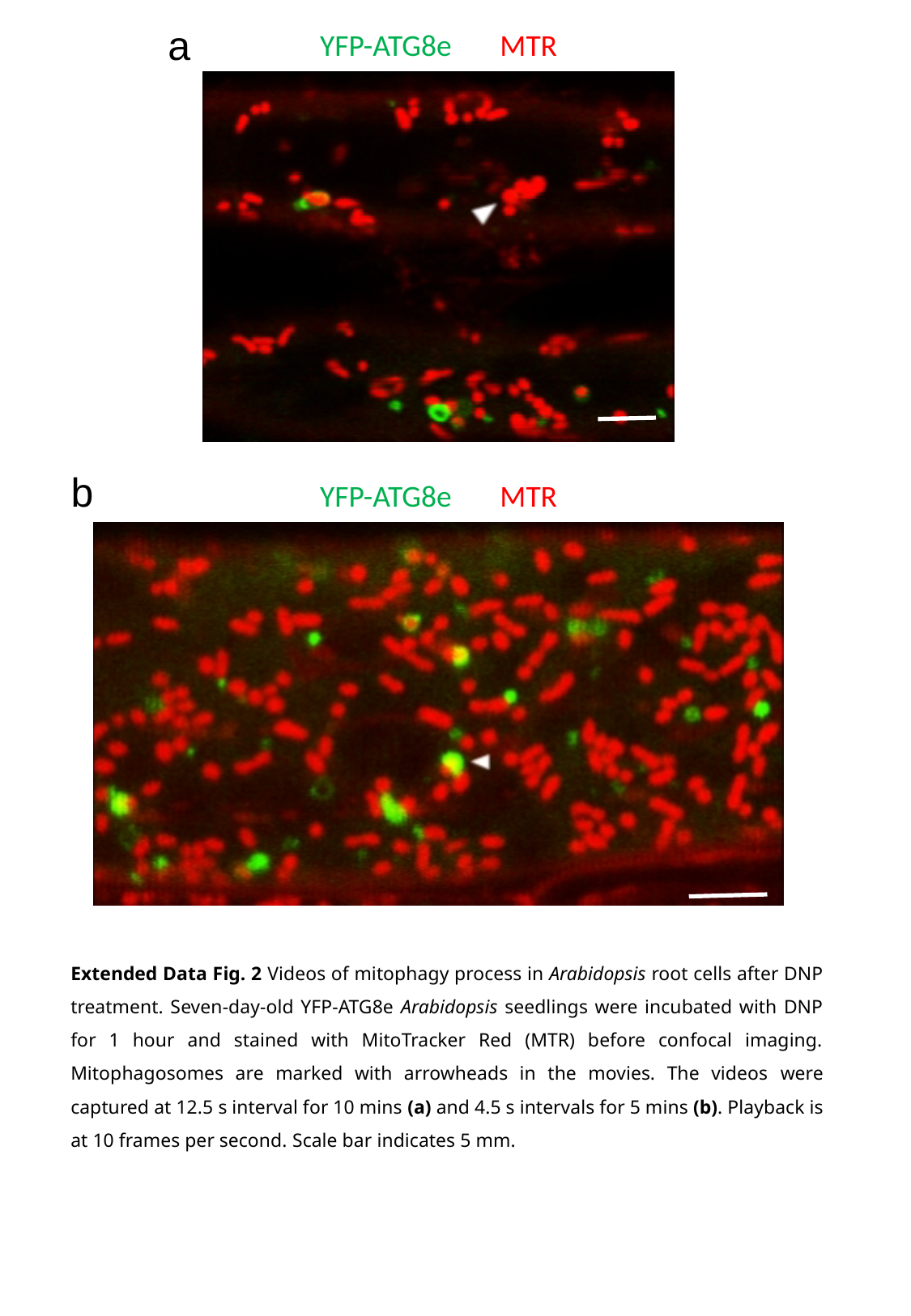

a
YFP-ATG8e MTR
b
YFP-ATG8e MTR
Extended Data Fig. 2 Videos of mitophagy process in Arabidopsis root cells after DNP treatment. Seven-day-old YFP-ATG8e Arabidopsis seedlings were incubated with DNP for 1 hour and stained with MitoTracker Red (MTR) before confocal imaging. Mitophagosomes are marked with arrowheads in the movies. The videos were captured at 12.5 s interval for 10 mins (a) and 4.5 s intervals for 5 mins (b). Playback is at 10 frames per second. Scale bar indicates 5 mm.

### Slide 3
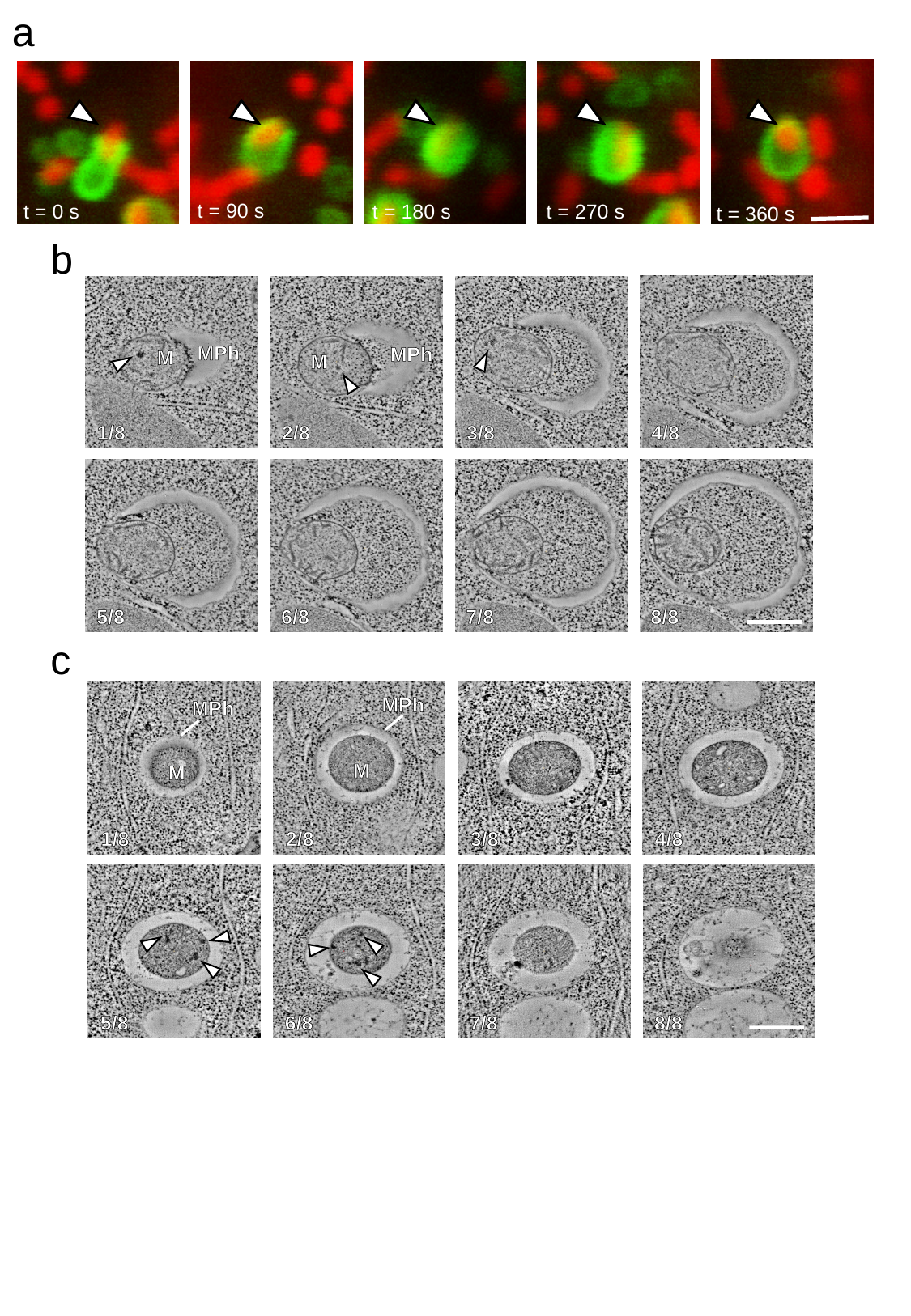

a
t = 90 s
t = 270 s
t = 180 s
t = 0 s
t = 360 s
b
MPh
MPh
M
M
1/8
2/8
3/8
4/8
5/8
6/8
7/8
8/8
c
MPh
MPh
M
M
1/8
2/8
3/8
4/8
5/8
6/8
7/8
8/8

### Slide 4
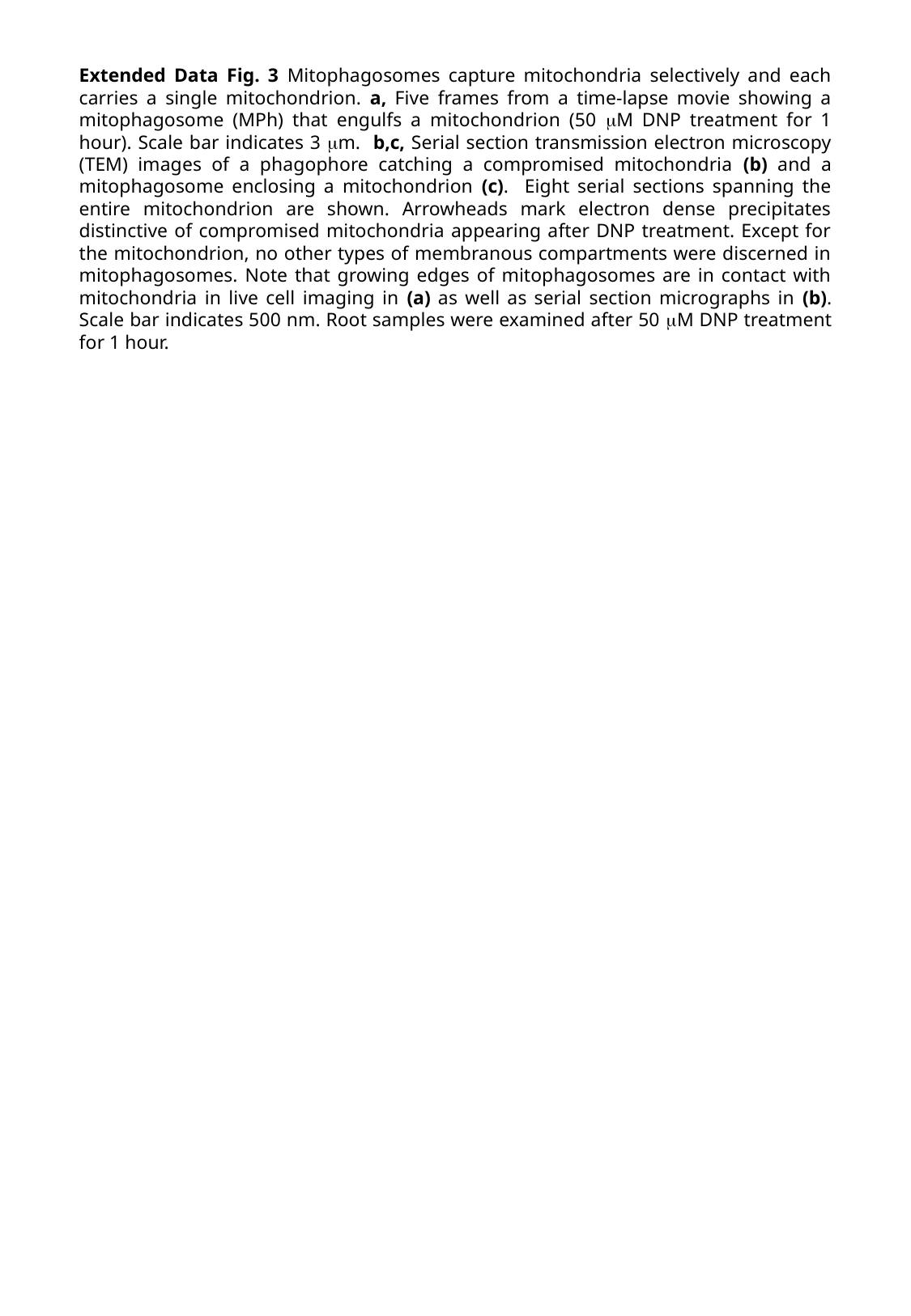

Extended Data Fig. 3 Mitophagosomes capture mitochondria selectively and each carries a single mitochondrion. a, Five frames from a time-lapse movie showing a mitophagosome (MPh) that engulfs a mitochondrion (50 μM DNP treatment for 1 hour). Scale bar indicates 3 μm. b,c, Serial section transmission electron microscopy (TEM) images of a phagophore catching a compromised mitochondria (b) and a mitophagosome enclosing a mitochondrion (c). Eight serial sections spanning the entire mitochondrion are shown. Arrowheads mark electron dense precipitates distinctive of compromised mitochondria appearing after DNP treatment. Except for the mitochondrion, no other types of membranous compartments were discerned in mitophagosomes. Note that growing edges of mitophagosomes are in contact with mitochondria in live cell imaging in (a) as well as serial section micrographs in (b). Scale bar indicates 500 nm. Root samples were examined after 50 μM DNP treatment for 1 hour.

### Slide 5
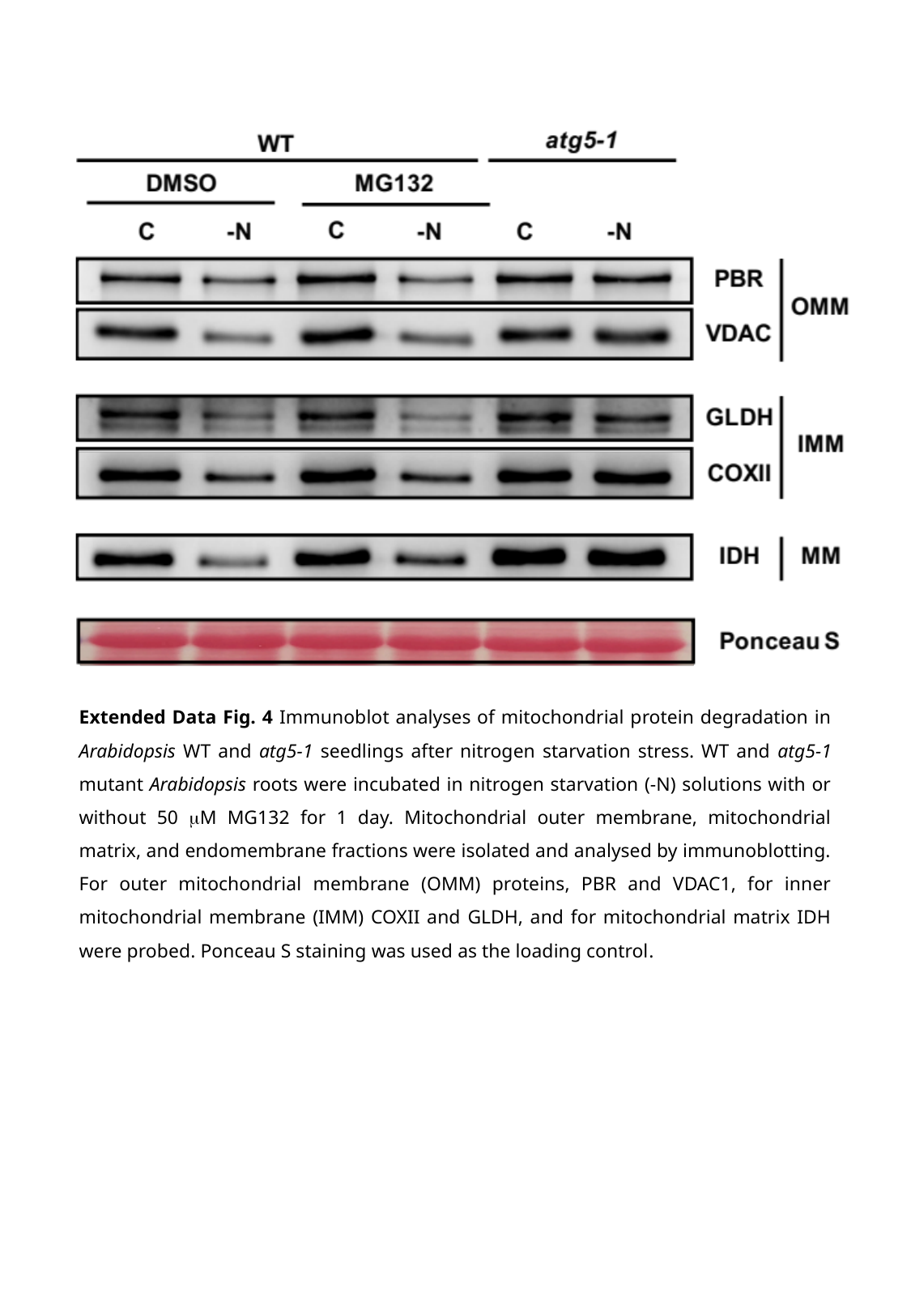

Extended Data Fig. 4 Immunoblot analyses of mitochondrial protein degradation in Arabidopsis WT and atg5-1 seedlings after nitrogen starvation stress. WT and atg5-1 mutant Arabidopsis roots were incubated in nitrogen starvation (-N) solutions with or without 50 μM MG132 for 1 day. Mitochondrial outer membrane, mitochondrial matrix, and endomembrane fractions were isolated and analysed by immunoblotting. For outer mitochondrial membrane (OMM) proteins, PBR and VDAC1, for inner mitochondrial membrane (IMM) COXII and GLDH, and for mitochondrial matrix IDH were probed. Ponceau S staining was used as the loading control.

### Slide 6
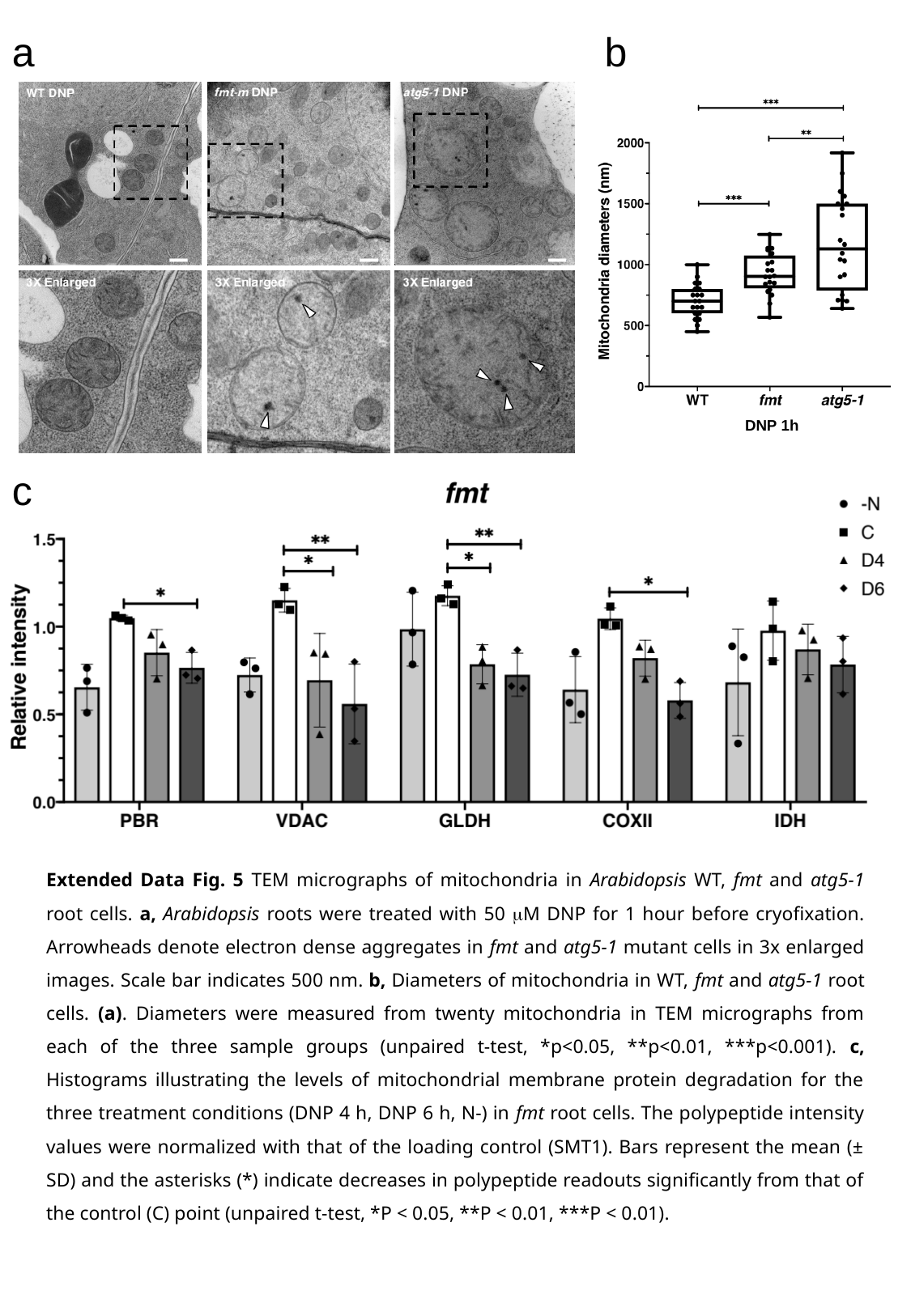

a
b
DNP 1h
c
Extended Data Fig. 5 TEM micrographs of mitochondria in Arabidopsis WT, fmt and atg5-1 root cells. a, Arabidopsis roots were treated with 50 μM DNP for 1 hour before cryofixation. Arrowheads denote electron dense aggregates in fmt and atg5-1 mutant cells in 3x enlarged images. Scale bar indicates 500 nm. b, Diameters of mitochondria in WT, fmt and atg5-1 root cells. (a). Diameters were measured from twenty mitochondria in TEM micrographs from each of the three sample groups (unpaired t-test, *p<0.05, **p<0.01, ***p<0.001). c, Histograms illustrating the levels of mitochondrial membrane protein degradation for the three treatment conditions (DNP 4 h, DNP 6 h, N-) in fmt root cells. The polypeptide intensity values were normalized with that of the loading control (SMT1). Bars represent the mean (± SD) and the asterisks (*) indicate decreases in polypeptide readouts significantly from that of the control (C) point (unpaired t-test, *P < 0.05, **P < 0.01, ***P < 0.01).
